# *In vivo* tissue-specific chromatin profiling in *Drosophila melanogaster* using GFP-tagged nuclei

**DOI:** 10.1101/2021.03.23.436625

**Authors:** Juan Jauregui-Lozano, Kimaya Bakhle, Vikki M. Weake

## Abstract

The chromatin landscape defines cellular identity in multicellular organisms with unique patterns of DNA accessibility and histone marks decorating the genome of each cell type. Thus, profiling the chromatin state of different cell types in an intact organism under disease or physiological conditions can provide insight into how chromatin regulates cell homeostasis *in vivo*. To overcome the many challenges associated with characterizing chromatin state in specific cell types, we developed an improved approach to isolate *Drosophila* nuclei tagged with GFP expressed under Gal4/UAS control. Using this protocol, we profiled chromatin accessibility using Omni-ATAC, and examined the distribution of histone marks using ChIP-seq and CUT&Tag in adult photoreceptor neurons. We show that the chromatin landscape of photoreceptors reflects the transcriptional state of these cells, demonstrating the quality and reproducibility of our approach for profiling the transcriptome and epigenome of specific cell types in *Drosophila*.

## Introduction

Dynamic regulation of the epigenome is crucial to replication, transcription, and DNA repair. For instance, accessible chromatin is associated with gene regulatory sequences, such as enhancers, promoters and transcription factor binding sites, and contributes to transcription initiation (Klemm et al., 2019). In addition, chromatin-associated proteins, such as histones, transcription factors or chromatin remodelers, modulate several processes, including nucleosome occupancy (Brahma & Henikoff, 2020), heterochromatin maintenance (Allshire & Madhani, 2018), and recruitment of DNA repair factors (Stadler & Richly, 2017). Thus, genome-wide chromatin profiling across different physiological states can help us understand how chromatin-mediated processes impact cell homeostasis.

The wide array of genetic manipulation tools, a highly mapped and annotated genome, relatively short lifespan, and ease of growth have made *Drosophila* one of the most widely used model organisms for studying the basic molecular mechanisms of eukaryotic cells (Hales et al., 2015). Further, the tissue homology between *Drosophila* and humans can be leveraged to uncover regulatory mechanisms associated with human relevant conditions, such as aging, neurodegeneration, and diabetes (Bolus et al., 2020; Graham & Pick, 2017; Piper & Partridge, 2018; Ugur et al., 2016). Since epigenetic dysregulation is one of the hallmarks of many diseases, including cancer and neurodegeneration (Bailey et al., 2018; Lardenoije et al., 2015), profiling chromatin states in a tissue-specific context using *Drosophila* might improve our understanding of how chromatin-associated changes contribute to disease onset. However, profiling cell type-specific chromatin states *in vivo* is challenging. Although tissue dissection can be coupled with bulk and single-cell genome wide experiments, manual tissue dissection is technically demanding and contamination from surrounding tissues can often confound results. To overcome these limitations, alternative techniques have been developed based around epitope labeling and immunoprecipitation of nuclei (Chitikova & Steiner, 2016). These nuclei tagging approaches, such as the “Isolation of Nuclei Tagged in specific Cell Types” (INTACT) method (Deal & Henikoff, 2010) have been applied to tissue specific experiments in *Arabidopsis* (Maher et al., 2018; Sijacic et al., 2018), *Drosophila* (Agrawal et al., 2019; Bozek et al., 2019; Henry et al., 2012; Jones et al., 2018), *Xenopus* (Amin et al., 2014), and mice (Ambati et al., 2016). In *Drosophila*, these nuclei labeling approaches rely on genetic tools for binary expression of transgenes, such as the well-established Gal4-UAS system (Brand & Perrimon, 1993). Currently, more than 8000 stocks that express Gal4 under control of different cell-type specific promoters are available through the Bloomington *Drosophila* Stock Center (BDSC). Thus, these nuclei tagging approaches combined with the Gal4-UAS expression system provide a powerful and flexible tool to manipulate and examine many cell-types in *Drosophila*.

We previously developed a Gal4-UAS based nuclei immuno-enrichment (NIE) protocol to isolate nuclei from specific *Drosophila* cell types labeled with an outer nuclear membrane localized GFP^KASH^ protein (Hall et al., 2017; Ma & Weake, 2014). This approach was successfully applied to transcriptomic studies in specific cell populations, such as larval glial cells (Ma et al., 2016), adult photoreceptor neurons (Hall et al., 2017, 2018), and olfactory sensory neurons (Slankster et al., 2020). However, our previous protocol yielded low nuclei numbers, which made performing chromatin profiling and obtaining material from rare cell populations challenging. Here, we sought to optimize the NIE protocol to increase nuclei yield and stringency over background. Using this ‘improved’ GFP^KASH^-based NIE protocol, we applied chromatin profiling techniques (Omni-ATAC, ChIP-seq and CUT&Tag) to NIE-purified adult *Drosophila* photoreceptor nuclei and demonstrate the reproducibility and quality of the associated datasets.

## Results

### Optimization of tissue-specific nuclei immuno-enrichment (NIE) from adult *Drosophila*

As a starting point for profiling chromatin states in specific cell types in *Drosophila*, we sought to improve nuclei yields obtained with the NIE protocol using flies that express the GFP^KASH^ tag in outer photoreceptor neurons driven by Rh1-Gal4 (herein referred as Rh1>GFP^KASH^) (Mollereau et al., 2000). We reasoned that isolating nuclei in a buffer designed to retain the integrity of the nuclear envelope would increase the availability of the GFP^KASH^ epitope, which is anchored to the outer nuclear membrane with GFP facing the cytoplasm (Fischer et al., 2004). Previous studies have shown that perinuclear proteins are retained when nuclei are purified using a detergent-containing isotonic buffer (Shaiken & Opekun, 2014), suggesting that the outer nuclear membrane remains intact under these conditions. Based on this rationale, we replaced the hypotonic/hypertonic buffers used in the homogenization, incubation, and washing steps of our previous NIE method with detergent-containing isotonic buffers. We also decreased the relatively high concentration of NP-40 detergent used for homogenization during the immunoprecipitation steps to decrease background binding (see methods). We refer to our previous and new NIE approaches as the ‘standard’ and ‘improved’ methods, respectively (Figure 1A).

**Figure 1.**
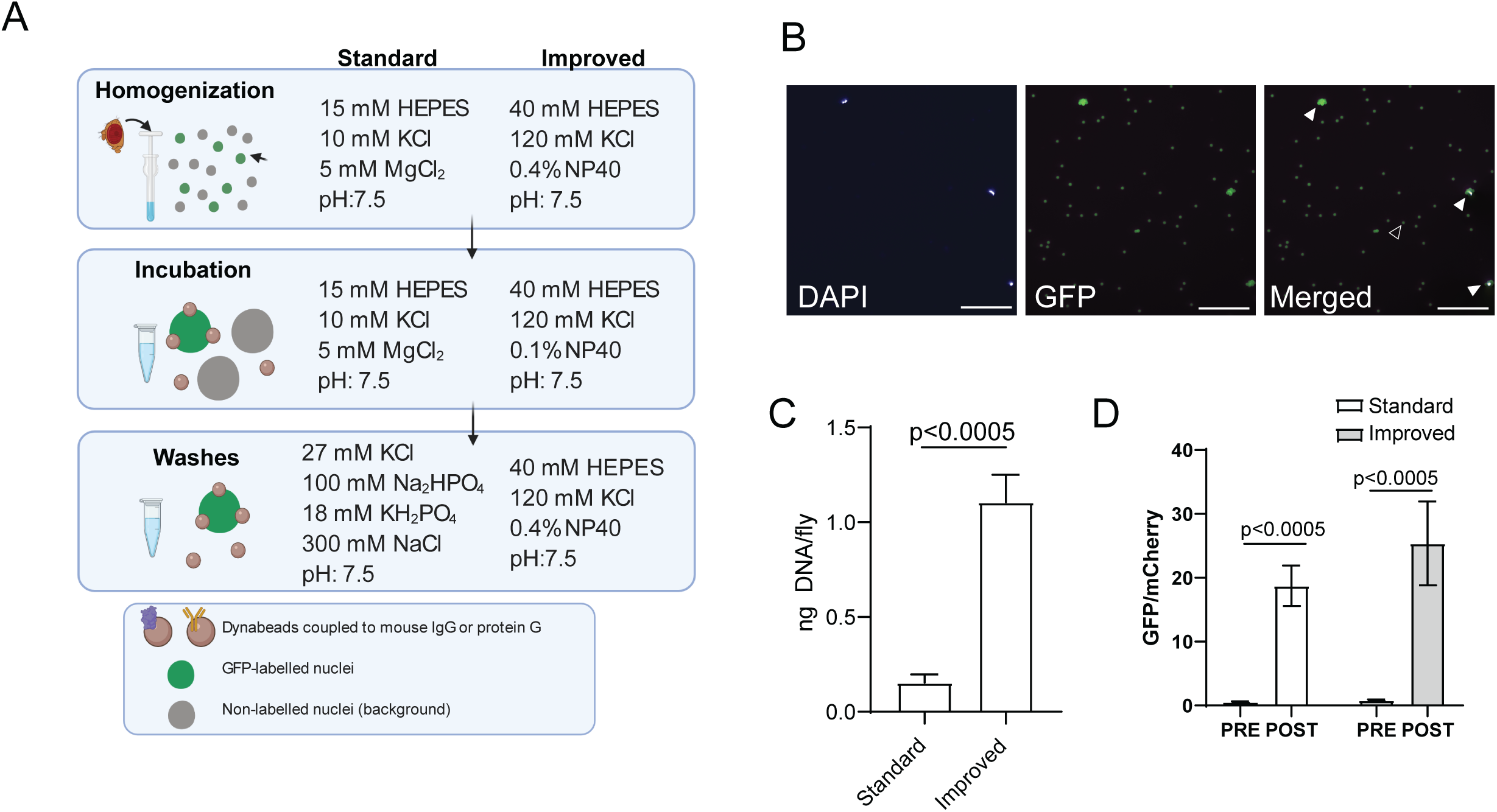
**A. Schematic diagram depicting the nuclear immuno-enrichment (NIE) protocol highlighting** major differences in buffer composition between the ‘standard’ and ‘improved’ methods. Heads from flies expressing Rh1>GFP^KASH^ were homogenized, followed by bead-antibody incubation and washes. B. Microscopy images of POST sample using the ‘improved’ method. Scale bars: 50 µM. White arrowhead: bead-bound nuclei. Black arrowhead: single bead. C. Bar plot showing DNA yields when Rh1>GFP^KASH^ nuclei were enriched using either the ‘standard’ or ‘improved’ NIE method (mean ± standard deviation (SD), n=4, p-value t-test). D. Bar plot showing qPCR enrichment for GFP and mCherry in the PRE and POST-NIE samples comparing ‘methods (mean ± SD; n=3, p-value t-test).

We first assessed how nuclei yields varied based on the NIE method used. To do this, we performed GFP^KASH^-based NIE using either the ‘standard’ or ‘improved’ method and quantified total DNA after each NIE reaction (n=4). We used DNA yield as a measure of nuclei yield because the magnetic beads used in the NIE auto fluoresce, making it difficult to quantify nuclei accurately using microscopy-based techniques (Figure 1B). The ‘improved’ method yielded 1.2 ng of DNA per fly, compared to 0.2 ng of DNA for the ‘standard’ method (Figure 1C). Considering that there are ∼7200 outer photoreceptors per fly, and that a diploid *Drosophila* nucleus typically contains ∼0.36 pg DNA (Rasch et al., 1971), the ‘improved’ method yields around 45% of the tagged nuclei compared with 13% for the ‘standard’ approach. We note that the starting material for each NIE reaction was 400 age-matched Rh1>GFP^KASH^ flies homozygous for both Gal4 and UAS transgenes; nuclei yield decreased approximately two-fold when GFP^KASH^-based NIE was performed using flies heterozygous for both transgenes (data not shown), suggesting that higher GFP^KASH^ expression levels can further improve purification efficiency.

Next, we evaluated if the NIE-purified nuclei were enriched relative to background cell types. To do this, we mixed an equivalent number of Rh1>GFP^KASH^ flies with Rh1>mCherry-FLAG^KASH^, performed GFP-based NIE, and extracted DNA before (PRE) and after (POST) immuno-enrichment. We then quantified the relative genomic copies of GFP and mCherry in each sample using quantitative PCR (qPCR). If nuclei from the POST sample are depleted of the mCherry^KASH^-positive nuclei upon GFP-based NIE, then the ratio of GFP/mCherry for the POST sample will be higher than the value of one observed in the PRE sample, which contains an equivalent number of GFP and mCherry labeled nuclei. Using this approach, we observed 24-fold enrichment of GFP nuclei over mCherry using the ‘improved’ method, which compared favorably with the 20-fold enrichment observed using the ‘standard’ method (Fig. 1C).

### Improved NIE method enriches for a purer cell-type specific nuclei pool relative to the standard

### method

Because we had previously generated high-quality nuclear RNA-seq from outer photoreceptor nuclei using the ‘standard’ approach (Hall et al., 2017), we profiled the nuclear transcriptome of NIE-purified outer photoreceptor nuclei (Rh1>GFP^KASH^) using the ‘improved’ method and compared the transcriptome between methods; we note that the identical genotype, sex, and age were used for both studies, and that both library sets were generated using the same amount of RNA. We first analyzed similarity between the two datasets by calculating Spearman correlation for gene counts (Figure 2A). Spearman’s rank scores between replicates were high for both methods (*p*<0.97), and samples clustered together based on the method used for NIE. Further, we also observed similar clustering by NIE approach using Principal Component Analysis (PCA) (Figure 2 – Supplemental Figure 1A). Notably, the variation between biological replicates slightly decreased using the improved method.

**Figure 2.**
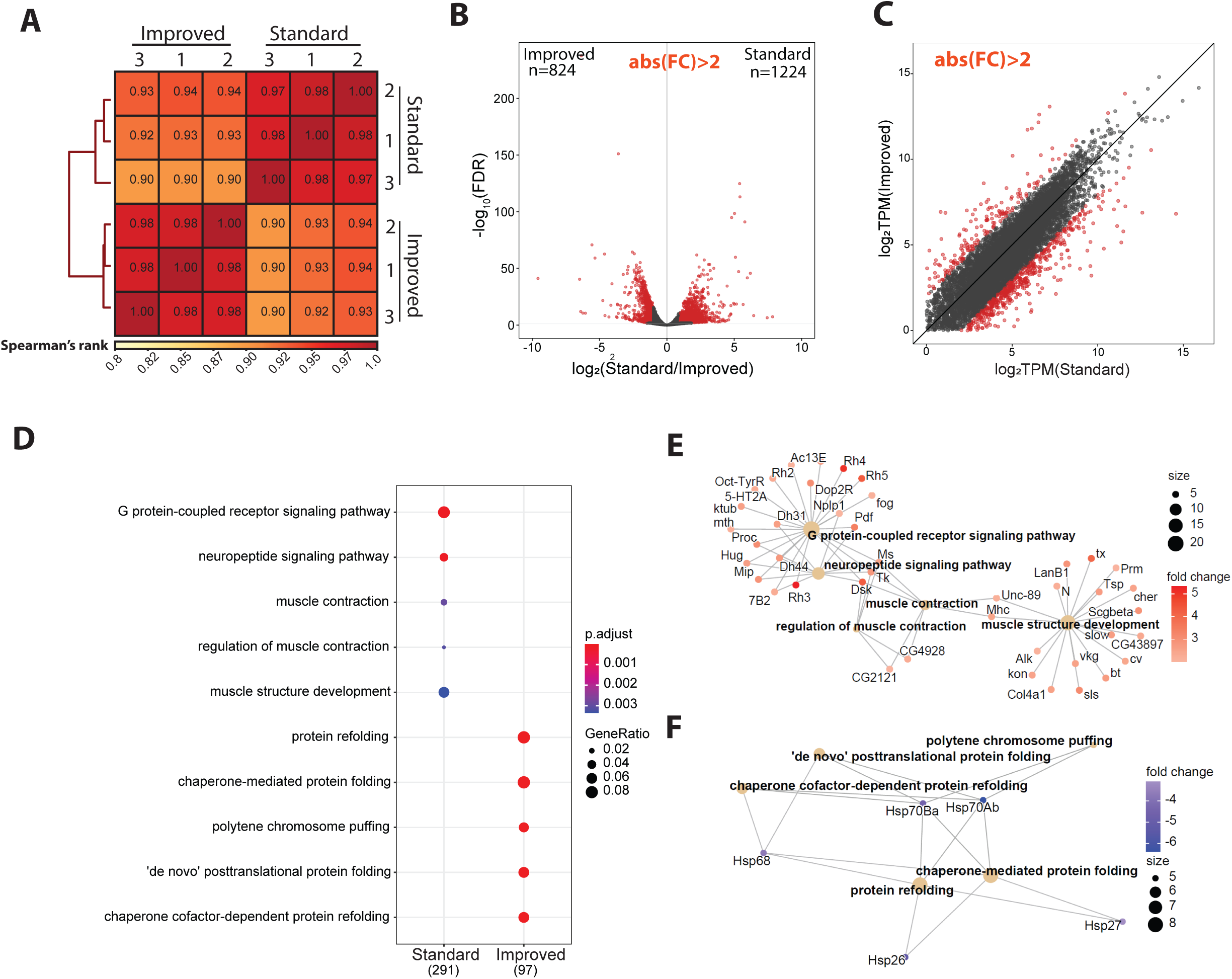
Improved NIE method enriches for a purer cell-type specific nuclei pool relative to the standard NIE method. A. Spearman correlation heatmap of gene expression profiles from nuclear RNA-seq of nuclei extracted with standard and improved method (n=4). Scores between 0 and 1 shown in each box correspond to Spearman’s rank score. B. Volcano plot showing the differentially expressed genes between methods. Genes with significant differential expression (FC > 2, FDR < 0.05) are highlighted in red. C. Scatter plot showing log2-transformed transcript per million (TPM) values between methods. DEGs highlighted in red, as in panel B. D. Gene Ontology (GO) term analysis on genes that are overrepresented in either the ‘standard’ or ‘improved’ method. E. Gene Concept Network plot (Cnetplot) highlighting linkage of individual genes and associated functional categories of genes over-represented in standard (top) and improved (bottom) dataset. Color intensity represents fold change between conditions.

The observation that samples clustered by method suggested there were differences between the datasets obtained using the different NIE methods. We sought to identify the differences in gene expression associated with each NIE method by analyzing differentially expressed genes (DEGs) (n=3). Surprisingly, we identified 2046 DEGs (FDR < 0.01, FC > 2) between the two NIE methods, despite their identical genotypes, sex, and age (Figure 2B). Amongst these genes, 824 genes were upregulated in the improved dataset, and 1224 genes were upregulated in the standard dataset, representing improved- or standard-enriched genes, respectively. RNA-seq libraries for each experiment were made using different RNA-seq kits (see methods). Since we used a kit designed for low-input material (200 pg – 10 ng RNA) to make the improved dataset libraries, we wondered if genes enriched in the improved dataset were being quantified as lowly-expressed in the standard dataset. However, the identified DEGs spanned a wide range of expression levels, including low, medium, and highly-expressed genes (Figure 2C), suggesting that differences in amplification of lowly abundant transcripts do not account for the differences in expression observed between the two approaches. Instead, inspection of the top DEGs in each condition revealed that several rhodopsin genes (*Rh3, Rh4,* and *Rh6*) were enriched in the standard method relative to the improved method. These rhodopsin proteins are highly enriched in inner photoreceptors (R7-R8) and are also expressed in the Johnston organ (Göpfert & Robert, 2001; Stark & Thomas, 2004), but are not expressed in outer photoreceptors; conversely, Rh1-Gal4 is expressed only in the outer photoreceptors (Mollereau et al., 2000). Since inner photoreceptor-specific genes were enriched in the standard dataset, these observations suggest that the ‘improved’ method yields a more tissue-specific enriched nuclei pool relative to our previous approach. GO-term analysis of genes that were upregulated in each dataset revealed that the standard-enriched DEGs were enriched for categories such as neuropeptide signaling pathway, muscle contraction, and muscle structure development (Figure 2D, top). Further, gene-concept network analysis revealed enrichment of 42 genes associated with non-photoreceptor cell types in the standard-enriched DEGs, including ventral lateral neuron-expressed *Pdf* (FBgn0023178), protocerebrum-enriched *Dsk* (FBgn0000500), and muscle-enriched *Unc-89* (FBgn0053519) (Figure 2E) (Helfrich-Förster & Homberg, 1993, Nichols et al., 1988). In contrast, GO terms over-represented in the improved-enriched DEGs were associated with processes related to protein folding and polytene chromosome puffing (Figure 2D, bottom). Gene-concept network analysis revealed that the over-representation of these GO-term categories were driven by a modest enrichment of five Heat Shock Protein (Hsp) genes (Figure 2F).

Altogether, these findings suggest that nuclei purified using the ‘improved’ NIE method have higher enrichment of tissue-specific transcripts compared to the ‘standard’ approach, which corresponds with the modest increase in GFP/mCherry ratio obtained in Figure 1D. Considering that the ‘improved’ method also had higher nuclei yields, we proceeded to optimize the subsequent chromatin profiling methods with NIE-purified outer photoreceptor nuclei from Rh1>GFP^KASH^ flies using this method.

### Profiling chromatin accessibility (Omni-ATAC) in NIE-purified nuclei

We next sought to profile accessible chromatin of NIE-purified nuclei using Omni-ATAC, a recently modified ATAC-seq technique which yields higher quality data, especially with lower input (Corces et al., 2017). ATAC-seq techniques, including Omni-ATAC, require optimization of the number of nuclei or cells used for each reaction to generate appropriate DNA fragment sizes and avoid either under- or over-tagmentation. Normally, cultured cells are counted to achieve precise numbers of cells per assay. However, nuclei bound to magnetic beads cannot be quantified using a cell counter because the free magnetic beads interfere with the identification of individual nuclei (see Figure 1B). To overcome this limitation, we isolated genomic DNA from a fraction of the purified nuclei and normalized input material for Omni-ATAC reactions based on this quantification (Figure 3A). We note that because our protocol begins with NIE-purified nuclei, mitochondria are already depleted from the initial starting material, as shown by qPCR analysis of mitochondrial DNA present in the PRE and POST NIE samples (Figure 3 - Supplemental Figure 1A). To evaluate whether differences in starting material would substantially alter data quality, we performed Omni-ATAC using either 50 or 100 ng of DNA (corresponding to approximately 125,000 and 250,000 nuclei, respectively) with a fixed amount of Tn5.

**Figure 3.**
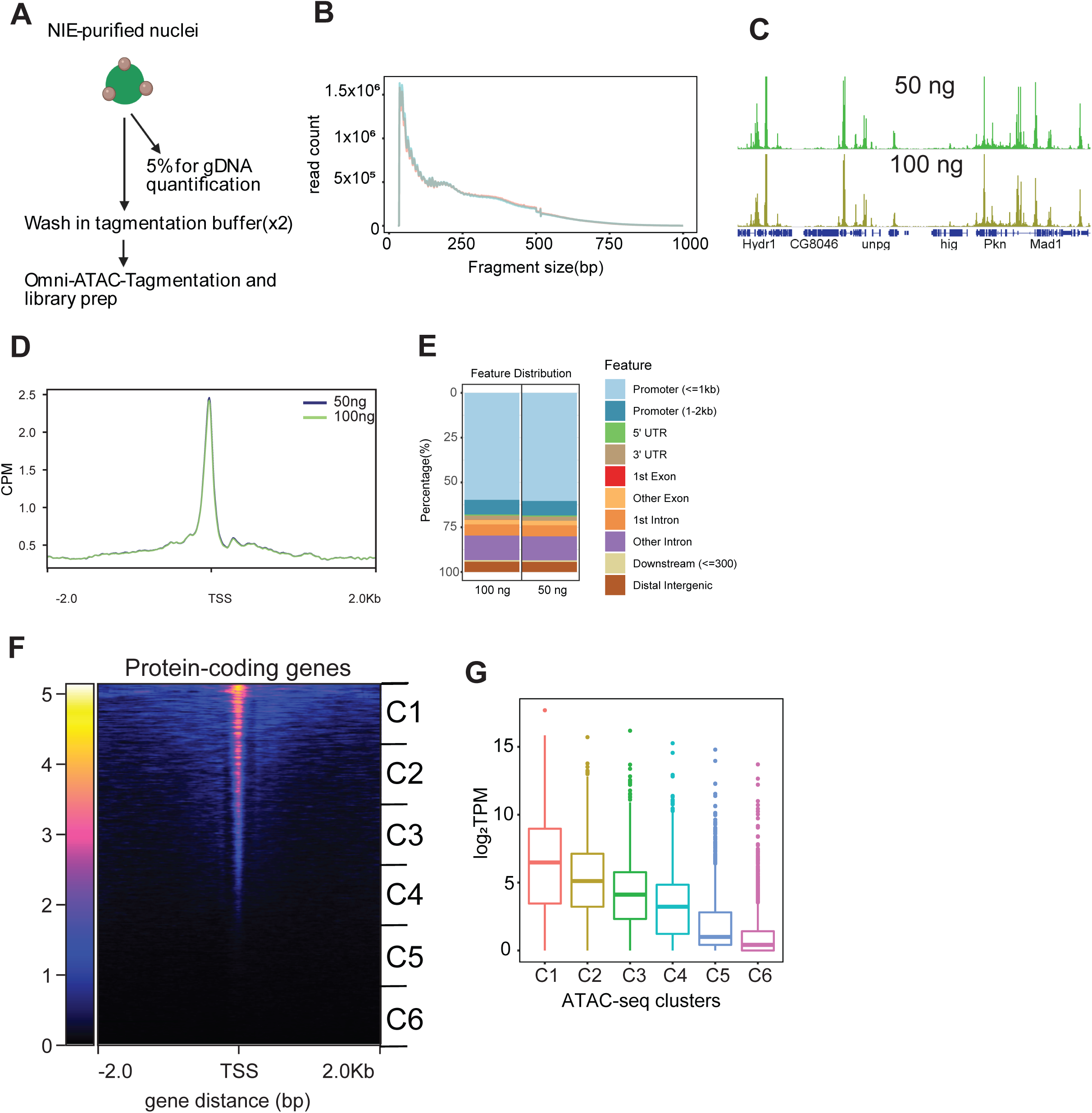
Profiling chromatin accessibility (Omni-ATAC) in NIE-purified nuclei. A. Diagram depicting Omni-ATAC approach applied to NIE-purified nuclei. After NIE purification, a fraction of nuclei is used for genomic DNA extraction and quantification to determine the input material for Omni-ATAC. Nuclei remain on ice until tagmentation, followed by two washes with tagmentation buffer without Tn5 enzyme. Upon washes, nuclei are tagmented using standard ATAC-seq conditions. B. Fragment size distribution of Omni-ATAC libraries using 50 ng (light green) or 100 ng (light red) as starting material. C. Genome browser views of counts per million (CPM)-normalized Omni-ATAC signal with genes shown in blue. D. Metaplot of CPM-normalized Omni-ATAC signal around the transcription start site (TSS) averaged for all protein-coding genes in the 50 ng and 100 ng samples. E. Genomic distribution of accessible peaks of 50 ng- and 100 ng- associated dataset. F. Heatmap showing CPM-normalized Omni-ATAC signal around TSS of protein-coding genes of 100ng-associated dataset. Clusters used for transcript boxplot are highlighted. G. Boxplot showing log_2_-transformed TPM scores for each cluster defined in 3F.

Tapestation analysis of Omni-ATAC libraries revealed similar DNA laddering patterns with both amounts of input nuclei (Figure 3 - Supplemental figure 1B). We then sequenced these libraries, and evaluated the size distribution of the mapped fragments. We observed the expected nucleosomal phasing distribution in both libraries (Figure 3B), with the first peak (80-120 bp) corresponding to nucleosome-depleted region (NDR)-associated DNA, followed by a peak around 180 bp corresponding to mononucleosome-associated fragments. Genome browser inspection of the data revealed discreet peaks with similar enrichment profiles obtained under each condition (Figure 3C). Since the Omni-ATAC signal should be enriched around transcriptional start sites (TSS), we next evaluated read distribution around the TSS of protein-coding genes (Figure 3D). We observed a significant enrichment of Omni-ATAC signal around the TSS with no differences between the 50 ng- and 100 ng- associated datasets. This finding was further corroborated by heatmap plots of all protein-coding genes ranked based on their Omni-ATAC signal enrichment around TSS (Figure 3 - Supplemental Figure 1C).

Next, we evaluated the genomic distribution of peaks from both samples (Figure 3E). As expected from the observed enrichment of Omni-ATAC signal around the TSS (Figure 3C), 70% of the peaks mapped to promoters with no discernible differences in distribution between the two samples. Because accessible chromatin is enriched for active promoters, we next evaluated if chromatin accessibility levels correlated with transcript levels detected by nuclear RNA-seq (see Figure 2). To do this, we divided the 13930 genes in the *Drosophila* genome based on their position on the heatmap into six groups, where genes are ranked based on the Omni-ATAC signal around the TSS (Figure 3F), and plotted the transcript level (log2 transcript per million - TPM) for all genes in each cluster (Figure 3G). We observed a positive correlation between the levels of chromatin accessibility at the TSS and transcript expression levels. Altogether, these observations suggest that high-quality chromatin accessibility data can be obtained from NIE-purified nuclei using as little as 50 ng of DNA equivalent of starting material, when coupled with Omni-ATAC.

### Omni-ATAC of NIE-purified nuclei does not require high sequencing depth

To benchmark the quality and reproducibility of the Omni-ATAC protocol using the NIE-purified nuclei, we sought to systematically evaluate different quality control metrics of ATAC-seq datasets. We performed Omni-ATAC on NIE-purified nuclei equivalent to 100 ng of DNA in four independent biological samples, processing and analyzing each replicate individually (n=4). We first calculated the Spearman’s correlation based on read distribution over a 500-bp binned genome, and found high reproducibility between samples, with Spearman’s *p* scores above 0.90 (Figure 4A). Next, we plotted the Omni-ATAC signal around the TSS of protein coding genes (Figure 4B). We observed that the enrichment profiles around the TSS were highly consistent between replicates, corroborating the Spearman’s correlation analysis. Next, we sought to evaluate the quality of peak-based analysis for each sample. Genome browser inspection of Omni-ATAC signal next to the peaks corresponding to each replicate showed high consistency, as determined by signal intensity of peaks (Figure 4C). Further, 88% of peaks presented significant overlap amongst all four replicates (Figure 4C). Similarly, we observed high concordance by Irreproducible Discovery Rate (IDR) analysis of peaks between replicates (Figure 4-Supplental Figure 1A), with all pair-wise comparisons having an IDR value above 0.61. The Fraction of Reads in Peaks (FRiP) score is a common quality control metric for genomic datasets, such as ChIP-seq and ATAC-seq, that measures overall signal-to-background ratio, as defined by ENCODE guidelines (Landt et al., 2012). According to ENCODE, good quality ATAC-seq datasets are defined as having FRiP score higher than 0.3. Thus, we next evaluated how FRiP scores varied based on sequencing depth. To do this, we down-sampled each replicate to 0.5, 1, 2.5, 5, 10, 20, 30, 40, and 50 million mapped fragments, and obtained its corresponding FRiP score (Figure 4E). FRiP scores did not vary significantly between replicates, and surprisingly, there was no substantial improvement in FRiP scores past 10 million mapped fragments. Further, visual inspection of the down-sampled data on a genome browser revealed similar enrichment of peaks at only 0.5 million fragments, resembling that observed using 50 million fragments (Figure 4 – Supplemental Figure 1B). Next, we evaluated the number of peaks called for each sample based on the number of fragments (Figure 4G). As expected, peak calling benefited from the higher sequencing depth. However, when the number of peaks identified was normalized to the sample with greatest sequencing depth (50 million mapped fragments), we found that obtaining 20 million fragments identified approximately 80% of all possible peaks. Taken together, these observations imply that Omni-ATAC datasets do not require high sequencing depth for consistent gene- and peak-based analysis, and that 10-20 million reads is likely sufficient for most peak-based analyses in *Drosophila* samples.

**Figure 4.**
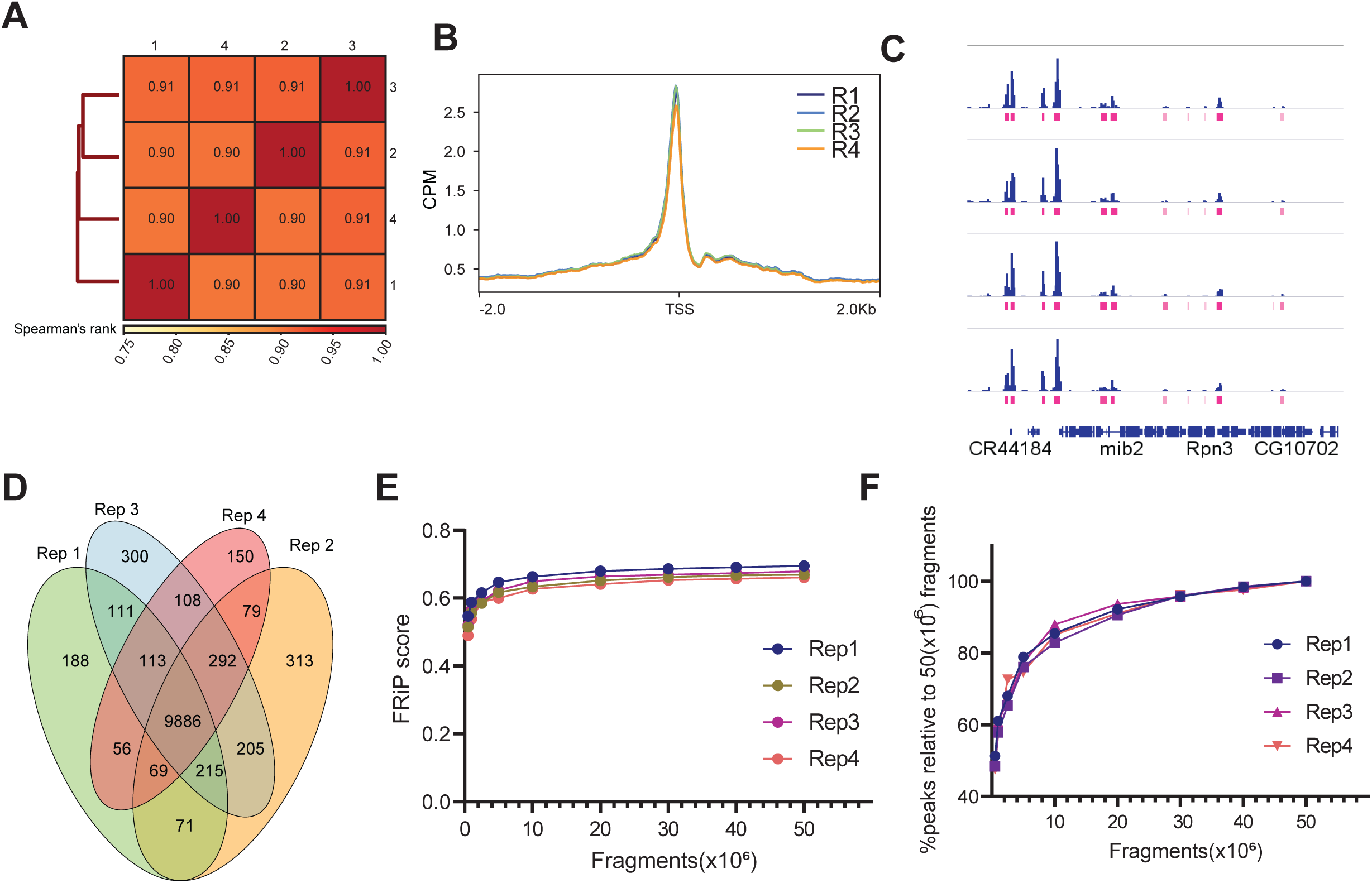
Omni-ATAC of NIE-purified nuclei does not require high sequencing depth. A. Spearman correlation heatmap of Omni-ATAC read distribution over binned genome. Scores between 0 and 1 shown in each box correspond to Spearman’s rank score. B. Metaplot of CPM-normalized Omni-ATAC signal around TSS averaged for all protein-coding genes comparing replicates (n=4). C. Genome browser inspection of CPM-normalized Omni-ATAC signal for each replicate, coupled with narrow peaks (pink). Genes are shown in blue. D. Venn diagram showing peak overlap/similarity between replicates. E. Fraction of Reads in Peaks (FRiP) scores of Omni-ATAC peaks comparing replicates down-sampled from 0.5 to 50 million mapped fragments. F. Percentage of peaks called relative to peaks called using the Omni-ATAC replicate #1, with 50×10^6^ mapped fragments as absolute percent of peaks.

### The histone methylation landscape of adult *Drosophila* photoreceptors

Chromatin Immunoprecipitation (ChIP) is one of the most commonly used techniques in the genomics field, whereby sonicated chromatin is used to immunopurify a protein-DNA complex, followed by purification of the enriched DNA. Coupled with qPCR or high-throughput sequencing (ChIP-seq), it allows researchers to interrogate if a protein of interest is bound to a particular locus, or assay its genome-wide distribution, respectively. We sought to optimize a ChIP protocol suitable for use with NIE-purified nuclei. During development of the protocol, we initially found that fixing the nuclei during homogenization led to an increase in background nuclei upon NIE (data not shown), leading us to modify the protocol so that the chromatin was cross-linked while the nuclei were immobilized on the magnetic beads, immediately following NIE (Figure 5A). Chromatin was then sonicated, and ChIP performed using standard approaches (see methods).

**Figure 5.**
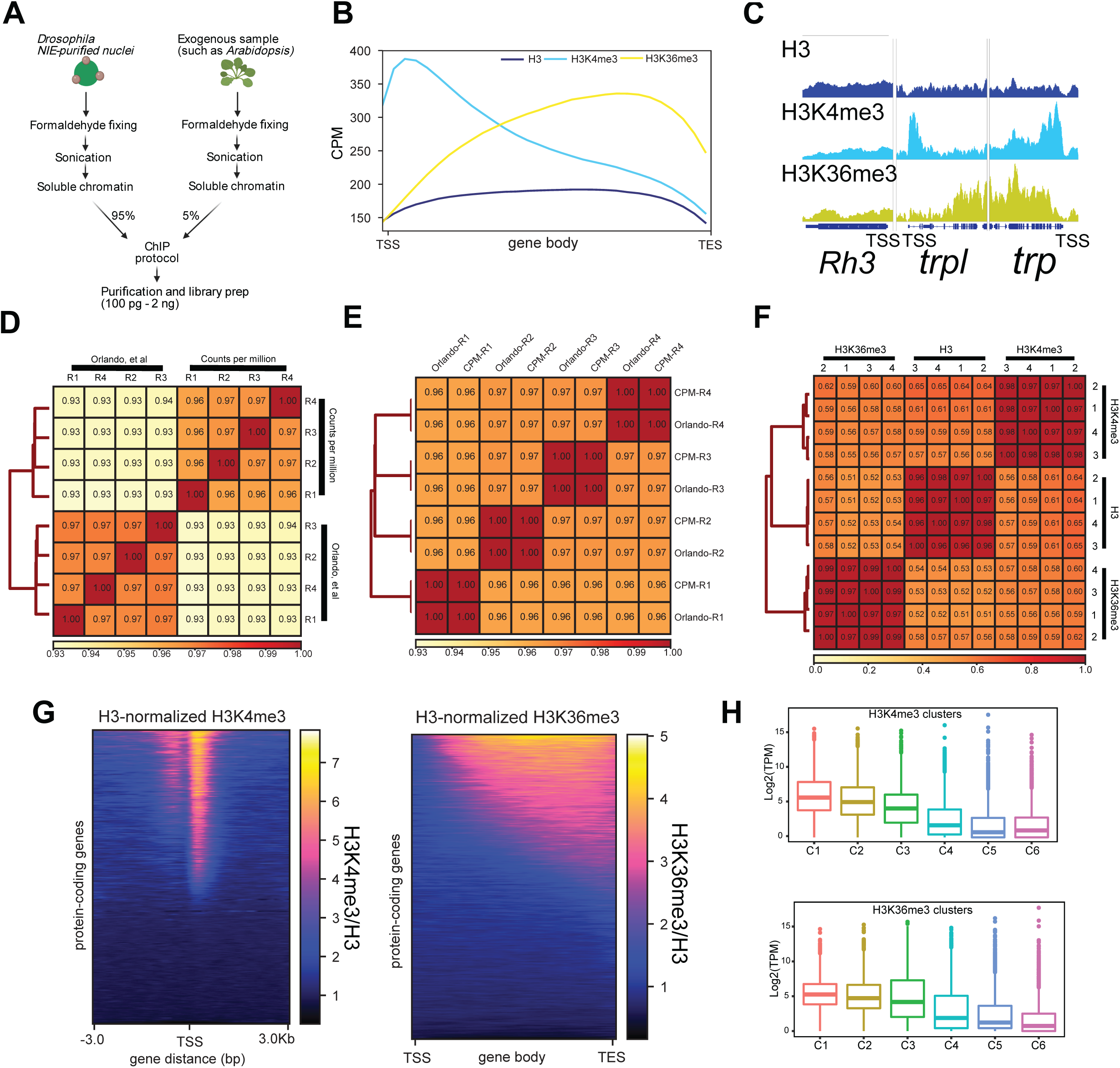
The histone methylation landscape of adult Drosophila photoreceptors. A. Diagram depicting Chromatin Immunoprecipitation (ChIP)-seq approach coupled to NIE-purified nuclei. Before adding the ChIP antibody, a fraction of soluble Drosophila chromatin (input) is quantified, to adjust final amount of chromatin per replicate, as well as to define amount of spike-in genome (In this case, 5% of Arabidopsis chromatin). B. Metaplots of H3 (dark blue), H3K4me3 (light blue) and H3K36me3 (yellow) ChIP-seq signal (CPM) over gene bodies averaged for all protein-coding genes. C. Genome browser inspection of H3, H3K4me3 and H3K36me3 ChIP-seq signal (CPM) around the inner photoreceptor-specific gene Rh3, which is not expressed in outer photoreceptors, and two highly expressed outer photoreceptor-specific genes trp and trpl. D. Spearman correlation heatmap of H3K4me3 ChIP-seq data comparing Spike-in and CPM normalization. Spearman’s rank scores are based on read distribution over binned genome. E. Spearman correlation heatmap of H3K36me3 ChIP-seq data comparing Spike-in and CPM normalization. Spearman’s rank scores are based on read distribution over binned genome. F. Spearman correlation heatmap of reads that align to binned genome for all replicates of H3, H3K4me3 and H3K36me3 ChIP-seq datasets. G. Heatmap showing signal for all protein coding genes of H3-normalized H3K4me3 (left) and H3-normalized H3K36me3 (right). F. Boxplots showing trans3c9ript level expressions of H3K4me3 (top) or H3K36me3 clusters (bottom).

To benchmark the ChIP protocol, we examined genome-wide distribution of two histone methyl marks, Histone H3 Lysine 4 tri-methylation (H3K4me3) and H3 Lysine 36 tri-methylation (H3K36me3), both of which have been widely characterized by ChIP-qPCR and ChIP-seq studies in *Drosophila* and other organisms. We also examined the distribution of bulk histone H3, as well as an input sonicated chromatin control. First, we assessed the enrichment of each antibody by evaluating the overall distribution of reads over gene bodies for all protein-coding genes. Histone H3 is distributed throughout both active and repressed chromatin, and is usually slightly depleted around the TSS of transcribed genes (Bai & Morozov, 2010). In *Drosophila,* as well as *in Saccharomyces cerevisiae* and in humans, H3K4me3 is enriched at the TSS whereas H3K36me3 localizes to gene bodies (Edmunds et al., 2008). Consistent with this expected distribution, we observed depletion of histone H3 and enrichment of H3K4me3 around the TSS, while H3K36me3 was enriched towards the 3’ region of the gene body (Figure 5B). Further, genome browser inspection of individual genes, such as the photoreceptor-enriched genes *trp* and *trpl,* corroborated the enrichment for H3K4me3 around the TSS and H3K36me3 over the gene body. In contrast, the inner photoreceptor-expressed *Rh3* showed no enrichment for either histone mark, as expected based on its lack of expression in outer photoreceptors (Figure 5C).

Next, we assessed the reproducibility between the replicates obtained using our ChIP-seq approach. Given the semi-quantitative nature of ChIP-seq, there has been growing interest in adding exogenous chromatin prior to immunoprecipitation, using the reads that map to the “reference” genome for spike-in normalization (Chen et al., 2016). To facilitate this spike-in normalization approach, we added 5% of *Arabidopsis thaliana* chromatin to *Drosophila* samples before each immunoprecipitation. To evaluate how the similarity between individual samples varied based on the normalization method, we normalized the data using the *Arabidopsis* spike-ins (as described in Orlando, et al.,2014) or calculated traditional counts per million or CPMs. We then calculated the Spearman correlation of read coverage over the binned genome for H3K4me3 and H3K36me3 separately (Figure 5D-E). Interestingly, the H3K4me3 samples clustered based on the normalization method used, although there were no major differences between Spearman’s rank scores obtained for individual samples using either approach. Replicate correlation was high for both normalization methods (*p* > 0.96 for both normalization methods). Strikingly, the H3K36me3 samples clustered together based on replicate rather than normalization approach, and each replicate had a *p=1,* with its normalization counterpart. Corroborating the heatmap findings, metaplot analysis of the H3K4me3 distribution around the TSS and H3K36me3 distribution over gene bodies showed no substantial differences between biological replicates using either normalization approach (Supplemental Figure 5A-B). To further assess similarity between the replicates based on antibodies used, we next evaluated Spearman’s correlation of CPM-normalized data for H3, H3K4me3, and H3K36me3 (Figure 5F). Corroborating the findings from the global read distribution over gene bodies, samples clustered together based on antibody. Moreover, the correlation between replicates for each antibody was also high (*p < 0.96).* Because H3K4me3 and H3K36me3 are histone modifications associated with active transcription, we next asked if H3K4me3 and H3K36me3 ChIP-seq signal levels positively correlated with gene expression. To do this, we ranked all protein-coding genes based on H3-normalized H3K4me3 signal around the TSS (Figure 5G, left) or H3-normalized H3K36me3 signal over gene bodies (Figure 5H, right), and separated all 13930 genes into six clusters based on their level of the respective histone mark. We then examined gene expression for each cluster by plotting transcript levels for each gene in the cluster (log2 transcript per million -TPM) (Figure 5H). Similar to our observations for the Omni-ATAC clusters, H3K4me3 and H3K36me3 levels positively correlated with active transcription.

Overall, these observations demonstrate that chromatin obtained from NIE-purified nuclei accurately reflect the transcriptional state of these cells and can be used for profiling of chromatin accessibility and histone modifications. Furthermore, in our hands, adding a reference genome for spike-in normalization does not outperform traditional CPM normalization. We note that although the ChIP-seq data shown here was generated from libraries that used 2 ng of DNA as starting material, libraries made with as little as 100 pg of DNA showed comparable profiles (Supplemental Figure 5C), suggesting that this ChIP-seq protocol is amenable to low-input starting material. We also performed qPCR on ChIP samples obtained using this protocol (Supplemental Figure 5D), demonstrating that this approach may be useful for researchers interested in examining individual genes rather than performing genome-wide studies.

### NIE-purified nuclei are compatible with CUT&Tag for profiling histone marks

Last, we sought to apply CUT&Tag to NIE-purified nuclei. CUT&Tag is a recently developed technique used to profile chromatin, whereby a fusion protein (pAG-Tn5) targets an antibody-bound chromatin target, followed by tagmentation and release of enriched DNA (Kaya-Okur et al., 2019). CUT&Tag has several advantages over ChIP-seq, including shorter sample processing times and lower background signal, therefore requiring less sequencing depth to identify high probability binding sites for proteins of interest. Further, CUT&Tag yields sequencing-ready libraries with no need for an additional library construction step. Based on these advantages, we sought to develop a CUT&Tag approach suitable for use with NIE-purified nuclei using commercially available Protein A/Protein G-Tn5 (pAG-Tn5).

Standard CUT&Tag protocols require cell/nuclei immobilization with Concanavalin A beads. However, NIE-purified nuclei are already bound to Protein G-magnetic Beads (PGBe), providing an initial starting point for CUT&Tag protocols. Our first H3K4me3 CUT&Tag trials with NIE-purified nuclei using PGBe were unsuccessful, and we wondered if the rabbit anti-H3K4me3 antibodies were being adsorbed by the excess protein G in our nuclei preparations (Figure 6A). To test this possibility, we performed NIE using Mouse IgG-coupled magnetic Beads (MIBe) instead of PGBe. Strikingly, performing NIE with MIBe led to successful purification of DNA following CUT&Tag, suggesting that PGBe were interfering with CUT&Tag steps. We then performed H3K4me3 CUT&Tag using age and sex-matched photoreceptor nuclei in order to compare the data with H3K4me3 ChIP-seq, since both datasets were obtained using the same antibody. TapeStation profiles of the four replicates detected sub-, mono- and di-nucleosomal fragments, with significant enrichment for mononucleosome-associated DNA (Figure 6B). We then proceeded with paired-end sequencing of the libraries. Genomic browser inspection of H3K4me3 CUT&Tag data (Figure 6C) revealed that profiles between replicates were highly consistent between the ChIP-seq and CUT&Tag methods. CUT&Tag enrichment is based on cleavage by Tn5, which traditionally binds and cuts accessible DNA. It has been shown that Tn5 can bind accessible chromatin during CUT&Tag, thereby increasing non-specific background. However, comparison of the CUT&Tag and Omni-ATAC profiles did not reveal substantial similarity, indicating that the CUT&Tag profiles obtained for H3K4me3 reflect the distribution of this mark rather than accessible chromatin. Next, we sought to systematically evaluate the signal to background ratio for CUT&Tag data relative to ChIP-seq. To do this, we down-sampled the H3K4me3 CUT&Tag and ChIP-seq samples to 0.5, 1, 2.5, 5, 10, and 15 million mapped fragments and calculated FRiP scores to assess quality of the data obtained using each approach (Figure 6D). Notably, CUT&Tag substantially outperformed ChIP-seq with a FRiP score of 0.367 for CUT&Tag data even at only 0.5 million mapped fragments. In comparison, the FRiP score for ChIP-seq data only reached 0.266 at 15 million fragments. However, analysis of the average H3K4me3 CUT&Tag signal around the TSS of all protein-coding genes revealed substantial differences between the individual replicates, both in intensity and distribution (Figure 6E, top). These differences were not observed for the ChIP-seq replicates (Figure 6E, bottom). Out of four biological replicates, only the metaplot profile of one CUT&Tag sample (replicate-4; R4) closely resembled the H3K4me3 ChIP-seq. To further assess the correlation between each CUT&Tag sample, we calculated Spearman’s correlation rank scores. Because CUT&Tag data had very low levels of background relative to ChIP-seq, we calculated the correlation based on read coverage over the narrow peaks obtained from the H3K4me3 ChIP-seq data (Figure 6G) instead of the binned genome. As expected from the above comparisons, samples clustered together based on technique. Using this approach, ChIP-seq samples had higher correlation values between individual replicates (*p*>0.9) compared with CUT&Tag replicates (*p*>0.83). R4(CUT&Tag) had the lowest correlation score when compared to the ChIP-seq samples, which is contrary to the profile obtained from the metaplot. To further assess if the same group of genes were being marked by H3K4me3 in both techniques, we ranked genes based on H3K4me3 ChIP-seq signal and compared the H3K4me3 CUT&TAG signal across replicates (Figure 6F). Heatmaps revealed that overall, CUT&Tag replicates showed similar patterns over the same group of genes. However, R4 had the highest similarity with ChIP-seq profiles in terms of overall distribution around the TSS.

**Figure 6.**
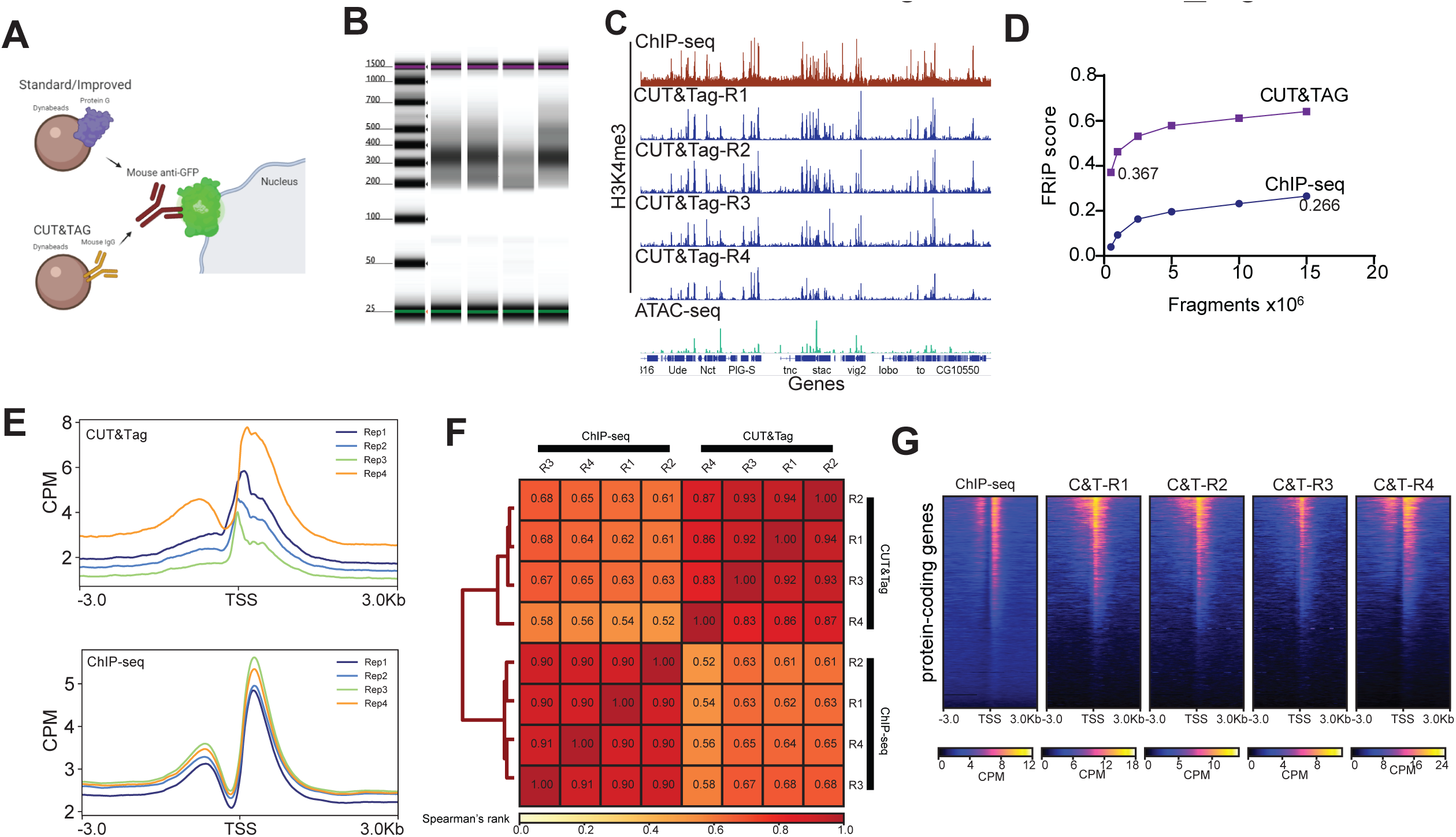
Bead modification in NIE protocol allows application of CUT&Tag. A. Schematic diagram representing the major difference between bead-antibody conjugation necessary to perform CUT&Tag in NIE-purified nuclei. Protein-G Dynabeads recognize both rabbit and mouse antibodies, while Mouse Pan IgG Dynabeads only recognize mouse antibodies. Nuclei preparation contains excess Dynabeads, therefore the protein G can interfere with CUT&Tag because it can bind the rabbit antibodies used to tag chromatin targets, such as H3K4me3. B. Tape Station profiles of H3K4me3 CUT&Tag libraries. C. Genome browser inspection (IGV) of CPM-normalized H3K4me3 ChIP-seq (top), H3K4me3 CUT&Tag replicates (medium) and Omni-ATAC (bottom). All samples were obtained from 10-day old male flies. Genes are shown in blue. D. FRiP score comparison between H3K4me3 CUT&Tag replicate 4 and H3K4me3 ChIP-seq replicate 1. Both samples were down sampled from 0.5 to 15 million mapped fragments. E. Metaplots of CPM-normalized H3K4me3 ChIP-seq (top) and H3K4me3 CUT&Tag (bottom) (n = 4 for each method). F. Heatmaps showing CPM-normalized H3K4me3 ChIP-seq (left-most) and H3K4me3 CUT&Tag signal for all replicates, with rows representing the same gene across all heatmaps. G. Spearman correlation heatmap of read distribution over H3K4me3 peaks called using ChIP-seq datasets. Correlation is calculated for H3K4me3 ChIP-seq and CUT&Tag replicates

Taken together, these observations indicate that a slight modification to the NIE reagents makes it possible to apply CUT&Tag to NIE-purified nuclei, providing a cost effective and efficient way of examining the genome-wide distribution of DNA-binding proteins. However, we note that the increased variability observed between CUT&Tag replicates relative to ChIP-seq samples suggests that further optimization to the protocol might improve reproducibility of these data for quantitative analysis.

## Discussion

Here, we demonstrate the feasibility of chromatin profiling in specific cell types using immuno-enriched nuclei as starting material and show that profiling of chromatin accessibility and histone modifications associated with active transcription correlate with the transcriptional state of the profiled cell type. Our NIE approach enables isolation of nuclei within one hour, that can be subsequently used for RNA, DNA, and chromatin extraction, therefore enabling the application of RNA-seq, ATAC-seq, ChIP-seq, and CUT&Tag (Figure 7A). By isolating nuclei, rather than cells, we can obtain highly pure nuclear RNA that provides a view of the actively transcribed genome. While these data correlate with the adult photoreceptor transcriptome determined in our previous studies using a similar approach (Hall et al., 2017), our modified NIE protocol results in significant decrease in levels of transcripts corresponding to genes that are expressed in other cell types. Thus, in addition to increasing nuclei yield, our improved NIE approach reduces levels of contamination from surrounding cells, with estimated purity levels of approximately 20-fold over background. Combining this improved NIE approach with library construction kits developed for low RNA inputs, such as the one used in this study, will facilitate RNA-seq studies on much rarer cell populations, or on cells labeled in mosaic animals, that have previously been difficult to analyze using other techniques.

**Figure 7.**
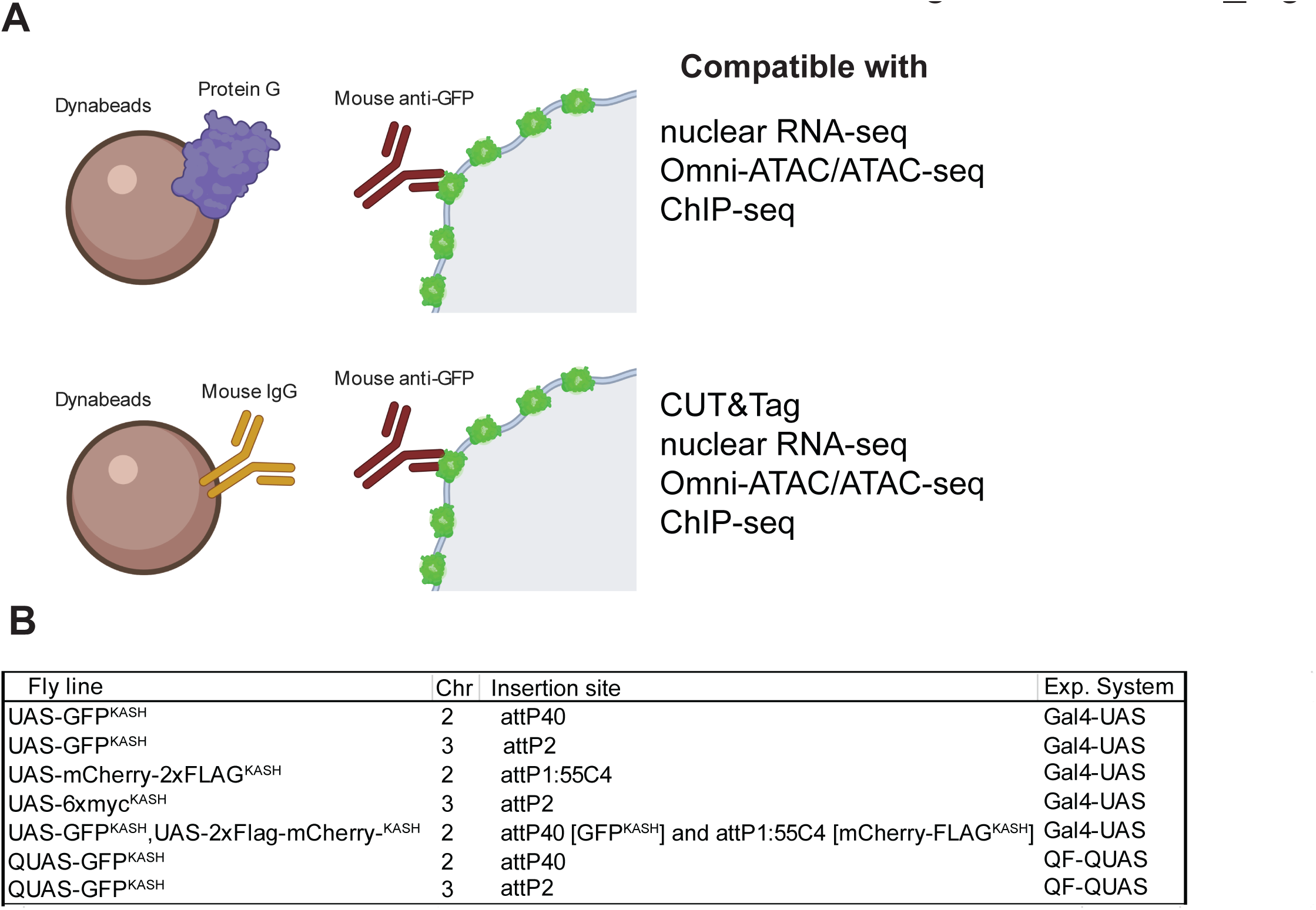
Method summary. A. Schematic diagram representing the two versions of the “improved” NEI-method. The first version (top) uses protein G-coupled magnetic Dynabeads, and can be coupled with RNA-seq, Omni-ATAC and ChIP-seq. The second version (bottom) uses Mouse IgG-coupled magnetic beads, and can be coupled with CUT&Tag, RNA-seq, Omni-ATAC and ChIP-seq. B. Table describing the available fly lines to perform NIE either using the Gal4-UAS or the QF-QUAS system.

In addition to RNA-seq, we profiled accessible chromatin at a genome-wide scale in the NIE-purified nuclei using Omni-ATAC. To our knowledge, this is the first report of cell-type specific chromatin accessibility data in adult *Drosophila*, although ATAC-seq studies have been performed in different embryonic cell-types isolated using the INTACT method (Bozek et al., 2019) and in dissected larval imaginal discs (Davie et al., 2015). Here, we show that using as little as 50 ng DNA equivalent of NIE-purified nuclei was sufficient to produce high-quality genome-wide chromatin accessibility data, suggesting that this technique should be suitable for lowly abundant cell types. Published reports have shown that ATAC-seq and Omni-ATAC can be applied to as little as 500 human cells (Buenrostro et al., 2013; Corces et al., 2017), indicating that these chromatin profiling approaches are highly amenable to low input samples.

We also applied two approaches to profile genome-wide distribution of histone modifications, ChIP-seq and CUT&Tag. Our ChIP-seq protocol is amenable to incorporation of exogenous chromatin for spike-in normalization, although in our hands, normalizing the ChIP-seq data with a published spike-in normalization approach did not outperform traditional CPM normalization. We note that there has been discussion of the caveats for spike-in normalization with regard to ChIP-seq data (refer to Dickson, et al., 2020). Last, we switched the beads used for NIE from protein-G Dynabeads to mouse IgG Dynabeads, allowing successful application of H3K4me3 CUT&Tag to NIE-purified nuclei. To our knowledge, this work is the first report of tissue-specific CUT&Tag in *Drosophila*. Although the CUT&Tag data showed increased variability between replicates relative to ChIP-seq, FRiP score evaluation showed that even at a low sequencing depth (1×10^6^ mapped fragments), H3K4me3 CUT&Tag signal-to-background ratio outperformed the ChIP-seq data obtained using the same antibody. We expect NIE-purified nuclei to be compatible with CUT&RUN techniques using a similar approach to that described in this study, since both techniques are based on the same principle; CUT&RUN uses MNase to digest and release enriched DNA (Skene & Henikoff, 2017).

Together, our data demonstrate that combining the improved NIE protocol with commonly used chromatin profiling techniques provides a feasible approach to characterizing the transcriptome and epigenome of specific cell types in *Drosophila*. Based on the wealth of available Gal4 drivers for cell type-specific expression in *Drosophila*, the NIE approach described here provides a flexible and resourceful chromatin profiling toolkit for researchers to interrogate chromatin-associated processes in a tissue-specific context. Additionally, we have generated fly stocks expressing the GFP^KASH^ tag under the Q binary expression system (Potter et al., 2010) as well as UAS lines that tag nuclei with either mCherry-FLAG, 6xmyc or mCherry-FLAG/GFP, to provide additional flexibility for studies in *Drosophila* (Figure 7B).

## Materials and Methods

### Fly strains

Flies homozygous for Rh1>GFP^KASH^ = *P{ry^+t7.2^=rh1-GAL4}3, ry^506^, P {*w*^+mC^ = UAS-GFP-Msp300KASH}attP2* or Rh1>mCherry^KASH^, *P{ry^+t7.2^=rh1-GAL4}3, ry^506^, P{w ^+mC^ = UAS-Msp300KASH-mCherry-Flag}attP2}* (Hall et al., 2017) were raised in 12:12 h light:dark cycle at 25°C on standard fly food. Flies were maintained in population cages with a density of ∼1000 flies/cage. Fresh food was switched every other day. For all the biological replicates, male flies were collected at 10 days post-eclosion at Zeitgeber time 6 (-/+ 1 hour).

### Nuclei Immuno-Enrichment (NIE)

NIE was performed as described previously (Hall et al., 2017; Ma & Weake, 2014) with minor modifications to the buffers used through-out the protocol. Briefly, fly heads from 400 age-matched flies were collected by freezing flies in 5 cycles of flash-freezing and vortexing. Fly heads were collected using frozen sieves and transferred to a 1 mL Dounce homogenizer containing 1 volume of homogenization buffer (40 mM HEPES, pH 7.5, 120 mM KCl and 0.4% v/v NP-40). Flies were homogenized using 10 strokes with ‘loose pestle’ followed by 10 strokes with ‘tight’ pestle. Homogenized lysate was then filtered using 40 μm cell strainers (Corning, Tewksbury MA, Catalog# 352340), and NP-40 was diluted to 0.1% final concentration by adding 3 volumes of Dilution buffer (40 mM HEPES, pH 7.5 and 120 mM KCl). Nuclei were immuno-enriched using 40 μL of Dynabeads Protein G (ThermoFisher, Waltham MA, Catalog #10004D) pre-coupled with 4 μg of mouse anti-GFP antibody (Sigma Aldrich, St. Louis MO, Catalog #11814460001) for RNA-seq, ChIP-seq and Omni-ATAC experiments. For CUT&Tag, nuclei were immunoenriched using 40 μL of Dynabeads Pan Mouse IgG (ThermoFisher. Catalog #11041) pre-coupled with 4 ug of mouse anti-GFP antibody (Sigma Aldrich, Catalog #11814460001). Beads and nuclei were incubated at 4°C for 30 min with constant rotation, followed by 3 x 5-min washes with homogenization buffer at 4°C.

### Quantitative PCR

DNA was purified with Quick-DNA Microprep Plus Kit (Zymo Research, Irvine CA, Catalog #D4074) and qPCR was performed using Bullseye EvaGreen qPCR 2X master mix-ROX (Midsci, Valley Park, MO, Catalog #BEQPCR-R) following manufacturer’s instructions.

### RNA-seq

Purified nuclei were resuspended in 100 μL TRI reagent (Zymo Research, Catalog #R2050-1-200). RNA was purified using Direct-zol™ RNA Microprep (Zymo Research, Catalog, #R2061) and quantified with Qubit™ RNA HS Assay Kit. 10 ng of nuclear RNA were used for construction of cDNA libraries with Ovation SoLo RNA-seq System with *Drosophila*-specific anyDeplete technology for rRNA depletion (Tecan, Redwood City, CA, Catalog #0502-32). Up to 16 libraries were pooled in one lane for paired-end 150 bp Illumina HiSeq sequencing.

### Omni-ATAC

Transposition was performed as published (Corces et al., 2017). Briefly, a fraction of immunoprecipitated nuclei were purified with Quick-DNA Microprep Plus kit (Zymo Research, Catalog #D4074). Nuclei corresponding to 50 or 100 ng were aliquoted and resuspended in 50 μL of Transposition mix (25 μL 2x TD buffer, 16.5 μL PBS, 0.05 μL 1% v/v Digitonin, 0.05 μL 10% v/v Tween and 2.5 μL TDE1 enzyme (Illumina, San Diego CA, Catalog #20034198). Tagmented DNA was purified with Zymo DNA clean & concentrator-5 kit (Zymo Research #D4013). Libraries were constructed using IDT for Illumina Nextera DNA Unique Dual Indexes Set A (Illumina, Catalog #20027213) and 7 PCR cycles were used to amplified libraries using NEBnext High-Fidelity 2X PCR Master Mix (New England Biolabs, Ipswich MA, Catalog #M0541S) and SYBR Green I (ThermoFisher, Catalog #S7563). To determine additional cycles, Nextera primers 1 and 2 were used. Purified libraries were submitted to a round of double-size selection with AMPure XP beads (Beckman Coulter, Brea CA, Catalog #A63880) with a 0.5X-1.0X ratio. Libraries fragment size distribution was assessed with TapeStation High-Sensitivity D1000 Screentapes (Agilent, Santa Clara CA, Catalog #5067-5584). Up to 16 libraries were pooled in one lane for paired-end 150 bp Illumina HiSeq sequencing.

### ChIP-seq

*Chromatin extraction (Drosophila):* Immunoenriched nuclei were resuspended in 1 mL of A1 buffer (15 mM HEPES, pH 7.5, 15 mM NaCl, 60 mM KCl, 4 mM MgCl2, 0.5% Triton X-100 v/v) and cross-linked with 1% methanol-free formaldehyde (ThermoFisher #28906) for 2 min at room temperature. Fixed nuclei were quenched with 125 mM Glycine, pH 7.5 for 5 min, followed by sonication in 130 μL of Nuclei Lysis Buffer (50 mM Tris-HCl, pH 8.0, 10 mM EDTA, 1% v/v SDS) in Covaris E220 with the following conditions: 10 min, 2% duty cycle, 105 Watts and 200 c.p.b. to obtain an average fragment size of ∼320 bp. Chromatin was centrifuged at 14,000 rpm, 10 min, 4°C, and the soluble chromatin supernatant was diluted with X-ChIP dilution buffer (16.7 mM Tris, pH 8.0, 167 mM NaCl, 1% Triton X-100 v/v, 1.2 mM EDTA pH 8.0), flash-frozen in liquid nitrogen, and stored at −20°C. *Chromatin extraction (Arabidopsis)*: 2.5 g of 10-day old *ref4-3MED15FLAG Arabidopsis* seedlings were ground to a fine powder using liquid nitrogen and resuspended in 20 mL of cold EB1 buffer (sucrose 0.440 mM, 10 mM Tris, pH 8.0, 10 mM MgCl2, 5 mM B-Me, 0.1 mM PMSF). The solution was filtered through two layers of miracloth and centrifuged at 3,000 x g, 20 min, 4°C. The pellet was then resuspended in 1 mL of cold EB2 Buffer (Sucrose 0.25M, 10 mM Tris, pH 8.0, 10 mM MgCl2, 1% v/v Triton X-100, 5 mM B-Me, 0.1 mM PMSF) and centrifuged at 4°C, 12,000 g for 10 min. The pellet was resuspended in 300 μL of cold EB3 buffer (sucrose 1.7M, 10 mM Tris, pH 8.0, 2 mM MgCl2, 0.15% v/v Triton X-100, 5 mM B-Me, 0.1 mM PMSF) and the sample was overlaid on top of 300 μL of cold EB3 and centrifuged at 4°C, 16,000g for 1 hour. Supernatant was transferred to a low-retention tube, snap-frozen and stored at −20°C. *Chromatin immunoprecipitation:* ChIP was performed as described (Deal & Henikoff, 2010) with the following modifications. Briefly, 380 ng of *Drosophila* chromatin (DNA) was mixed with 20 ng of *Arabidopsis* chromatin as a spike-in control (5%), and incubated with 1 μg of each of the following antibodies: H3 (Abcam, Cambridge MA, Catalog #1791), H3K4me3 (Abcam, Catalog #8580) and H3K36me3 (Abcam, Catalog #9050) for 12 to 18 hours at 4°C. Immunoprecipitated histone-DNA complexes were incubated with 25 μL Dynabeads protein G (ThermoFisher, Catalog #10004D) for 2 hours at 4°C, followed by 5-min washes with 1 mL Low Salt Buffer (20 mM Tris-HCl, pH 8.0, 150 mM NaCl, 0.1% v/v SDS, 1% v/v Triton X-100, 2 mM EDTA), 1 mL High Salt Buffer (20 mM Tris, pH 8.0, 500 mM NaCl, 0.1% v/v SDS, 1% v/v Triton X-100, 2 mM EDTA), 1 mL LiCl Wash buffer (10 mM Tris, pH 8.0, 250 mM LiCl, 0.1% v/v Na-Deoxycholate, 1% v/v NP-40 substitute, 1 mM EDTA) and 1 mL TE (10 mM Tris, pH 8.0, 1 mM EDTA). Histone-DNA complexes were eluted from the magnetic beads with X-ChIP elution buffer (100 mM NaHCO3, 1% v/v SDS), treated with RNAse A (ThermoFisher, Catalog #EN0531) at 37°C for 30 min and Proteinase K (ThermoFisher, Catalog #AM2546) at 55°C for 12 to 18 hours. DNA was purified with Zymo Research ChIP DNA clean & concentrator kit (Zymo Research, Catalog #D5205). Purified DNA was quantified with Qubit 1X HS DNA kit (ThermoFisher, Catalog #Q33230). Input sample fragment size was determined with TapeStation High-Sensitivity D5000 Screen tapes (Agilent, Catalog #5067-5592)

*ChIP-seq library prepation*: 2 ng of DNA were used for ChIP-seq libraries constructed with Tecan Ovation Ultralow V2 DNA-Seq Library Preparation Kit-Unique Dual Indexes (Tecan, Catalog #9149-A01). Following amplification, purified libraries were submitted to a round of double-size selection with AMPure XP beads (Beckman Coulter, Catalog# A63880) with a 0.61X-0.8X ratio. Libraries fragment size distribution was assessed with TapeStation High-Sensitivity D1000 Screentapes (Agilent, Catalog #5067-5584). Up to 16 libraries were pooled in one lane for paired-end 150 bp Illumina HiSeq sequencing.

### CUT&Tag

CUT&Tag was performed using CUTANA™ CUT&Tag reagents (Epicypher, Durham NC, #15-1017, #15-1018, #13-0047) following manufacturer’s “Direct-to-PCR CUT&Tag Protocol” with minor modifications: Briefly, purified nuclei were washed 3 times with cold Antibody150 buffer, and protocol was started at Section III “Binding of Primary and Secondary antibodies” and followed as described: https://www.epicypher.com/content/documents/protocols/cutana-cut&tag-protocol.pdf

### Data processing

Raw reads were trimmed using Trimmomatic version 0.39 (Bolger et al., 2014) to filter out low quality reads (Q>30) and clean adapter reads. Cleaned reads were aligned to the *Drosophila melanogaster* genome (BDGP Release 6 + ISO1 MT/dm6 from UCSC) using splicing-aware aligner STAR version 1.3 (Dobin et al., 2013) for RNA-seq, and Bowtie2 version 2.3.5.1 (Langmead & Salzberg, 2012, p. 2) for Omni-ATAC, ChIP-seq and CUT&Tag using –sensitive settings. Samtools version 1.8 (Li et al., 2009) was used to obtain, sort and index BAM files. For genome browser inspection as well as further analyses, bigwig files were generated by normalizing datasets to count-per-million CPM coverage tracks using *deepTools* version 3.1.1 (Ramírez et al., 2014) using *--normalizeUsing CPM* settings. Spearman’s correlation scores were calculated using deepTools’ subpackages *multiBigwigSummary* and *plotCorrelation*. Metaplots and genomic distribution heatmaps were made with deepTools’ subpackages *computeMatrix, plotHeatmap* and *plotProfile*. GO term analysis was performed using R package *clusterProfiler* (Yu et al., 2012). *Spike-in normalization.* FastQ Screen version 0.13.0 (Wingett & Andrews, 2018) was used to separate reads that uniquely mapped to either the genome of *Drosophila melanogaster* (BDGP Release 6 + ISO1 MT/dm6 from UCSC) or *Arabidopsis thaliana* (Tair10 – Arabidopsis.org) using the filter option and with *sensitive* parameters. Each fastq file was aligned and processed separately, and alignment rates to each genome file were used to calculate spike-in factors (Orlando et al., 2014). Calculated spike-in factors were used to convert bam files into normalized bigwig files using deepTools *bamCoverage* subpackage, with *–scaleFactor* setting, generating Reference-adjusted Reads Per Million (RRPM) files with a 10-bp resolution. Encode blacklist regions were removed. Spearman correlation scores were calculated by partitioning the mappable genome into 500-bp bins and obtaining the RRPM within each bin. Omni-ATAC narrow peaks were obtained using MACS2 version 2.1.2 (Zhang et al., 2008) with settings: “--nolambda --nomodel --extsize 150 --shift 75 --keep-dup all”, and H3K4me3 ChIP-seq and CUT&Tag peaks were obtained with settings: “--nolambda --nomodel -- keep-dup all”. FRiP scores were calculated using *FeatureCounts* of Subread version 1.6.1. (Liao et al., 2013). Peak overlap and genomic distribution of peaks was determined using R package ChIPseeker (Yu et al., 2015).

### Graph plots

Bar-plots were generated using GraphPad Prism and scripts used for RNA-seq analysis and plot generation are available upon request.

## Acknowledgements

We thank all members of the Weake lab and Dr. Hana Hall for their suggestions for the manuscript. We also thank Dr. Ulrike Litzenburger for her assistance during Omni-ATAC troubleshooting, and Dr. Xiangying (Candy) Mao and Dr. Clint Chapple for providing the *Arabidopsis thaliana* samples. Information from FlyBase was used in this study. Support from the American Cancer Society Institutional Research Grant (IRG #58-006-53) to the Purdue University Center for Cancer Research is gratefully acknowledged. The RNA-seq work was supported, in part by the Indiana Clinical and Translational Sciences Institute funded by Award Number UL1TR002529 from the National Institutes of Health, National Center for Advancing Translational Sciences, Clinical and Translational Sciences Award. Research reported in this publication was also supported by a Bird Stair Research Fellowship (Biochemistry Department, Purdue University) to JJL, and by the National Eye Institute of the NIH under Award Number R01EY024905 to VMW.

## Data Availability

Previously published RNA-seq expression data are accessible through Gene Expression Omnibus (GEO) repository under series accession number GSE83431. Data obtained for this manuscript are accessible to reviewers through GEO repository under series accession number GSE169328

---

**Figure 2-Supplemental Figure 1.**
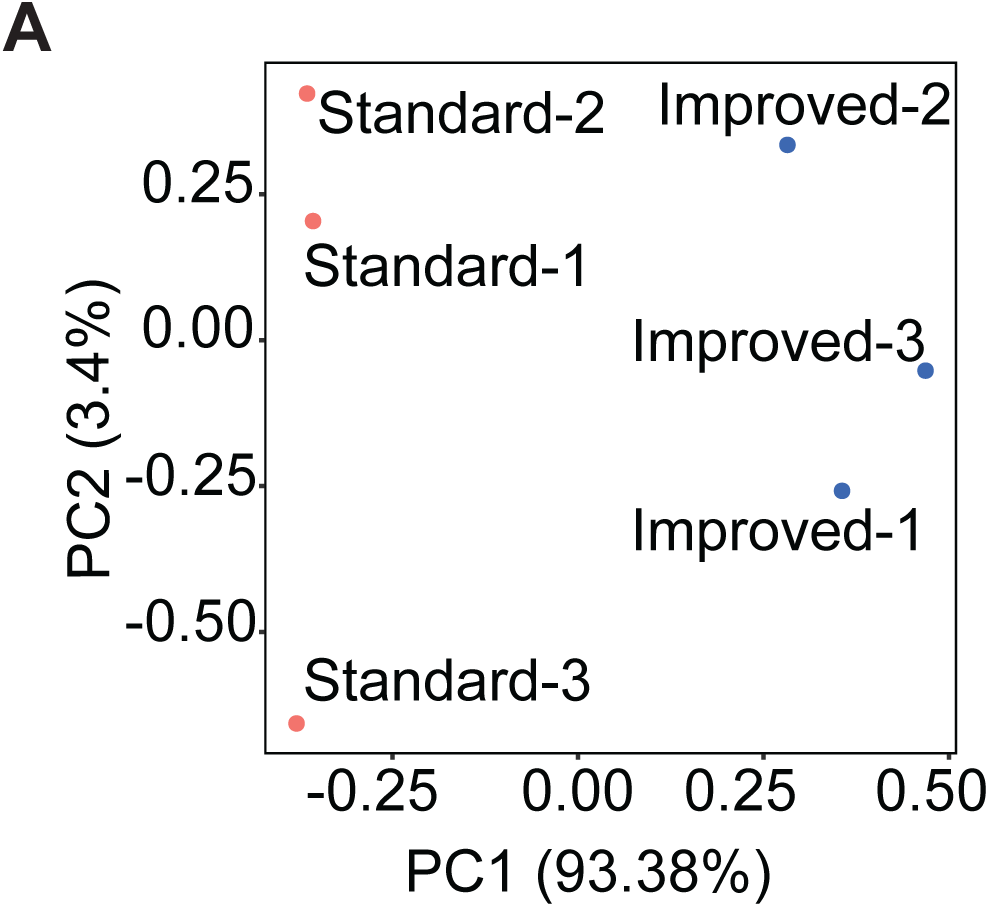
A. Principal Component Analysis (PCA) of gene counts

**Figure 3-Supplemental Figure 1.**
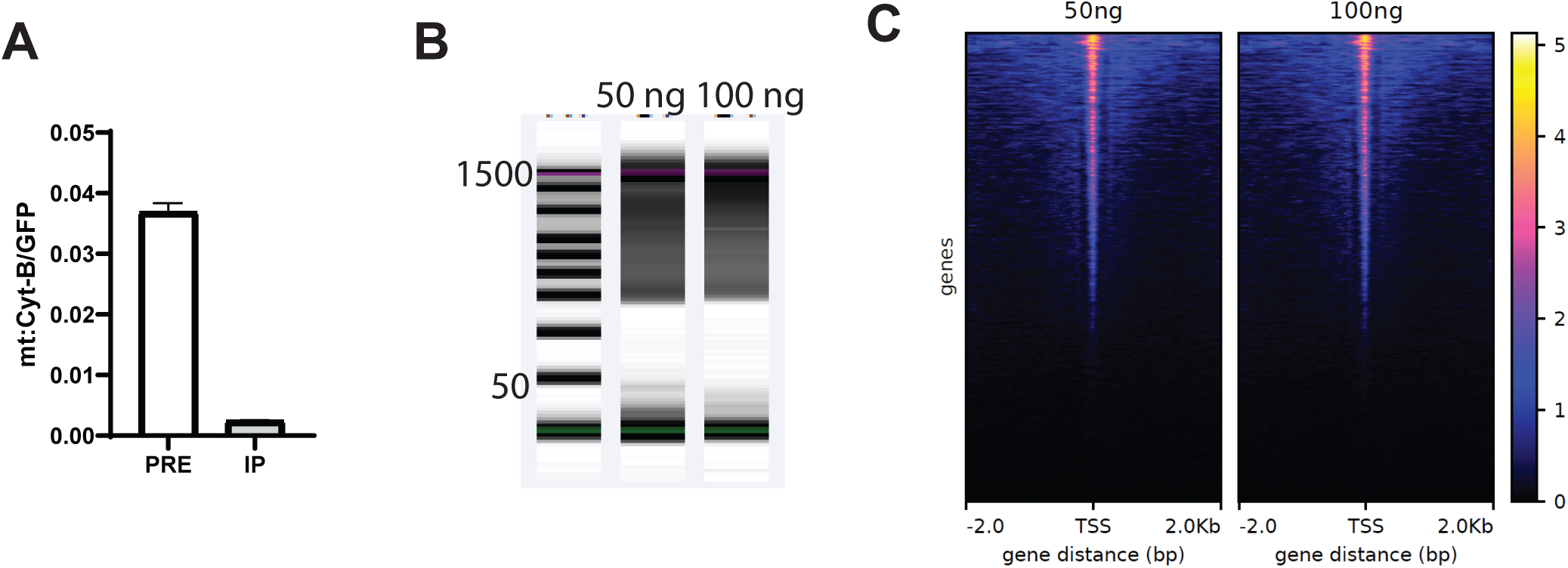
A. Bar plot showing qPCR enrichment for GFP and mitochondrial DNA (mt:Cyt-B) in the PRE and POST-NIE. (Mean ± SD; n=3). B. Tapestation profiles of Omni-ATAC libraries prepared using 50 ng (Blue) and 100 ng (Orange) datasets. C. Heatmaps showing CPM-normalized Omni-ATAC signal for 50 ng- and 100 ng-associated datasets.

**Figure 4-Supplemental Figure 1.**
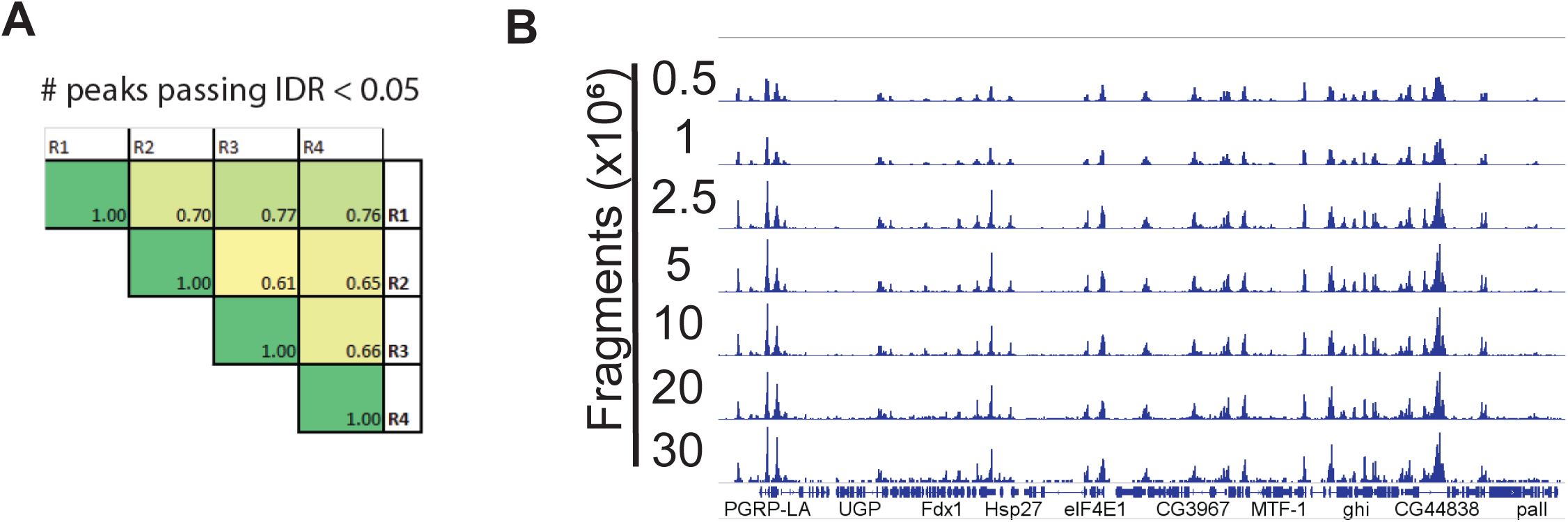
A. Pair-wise comparison of irreproducible discovery rate (IDR) values of peaks that pass the 0.05 threshold. B. Genome browser inspection of down-sampled CPM-normalized Omni-ATAC signal used for FRiP score analysis. Genes are shown in blue.

**Figure 5-Supplemental Figure 1.**
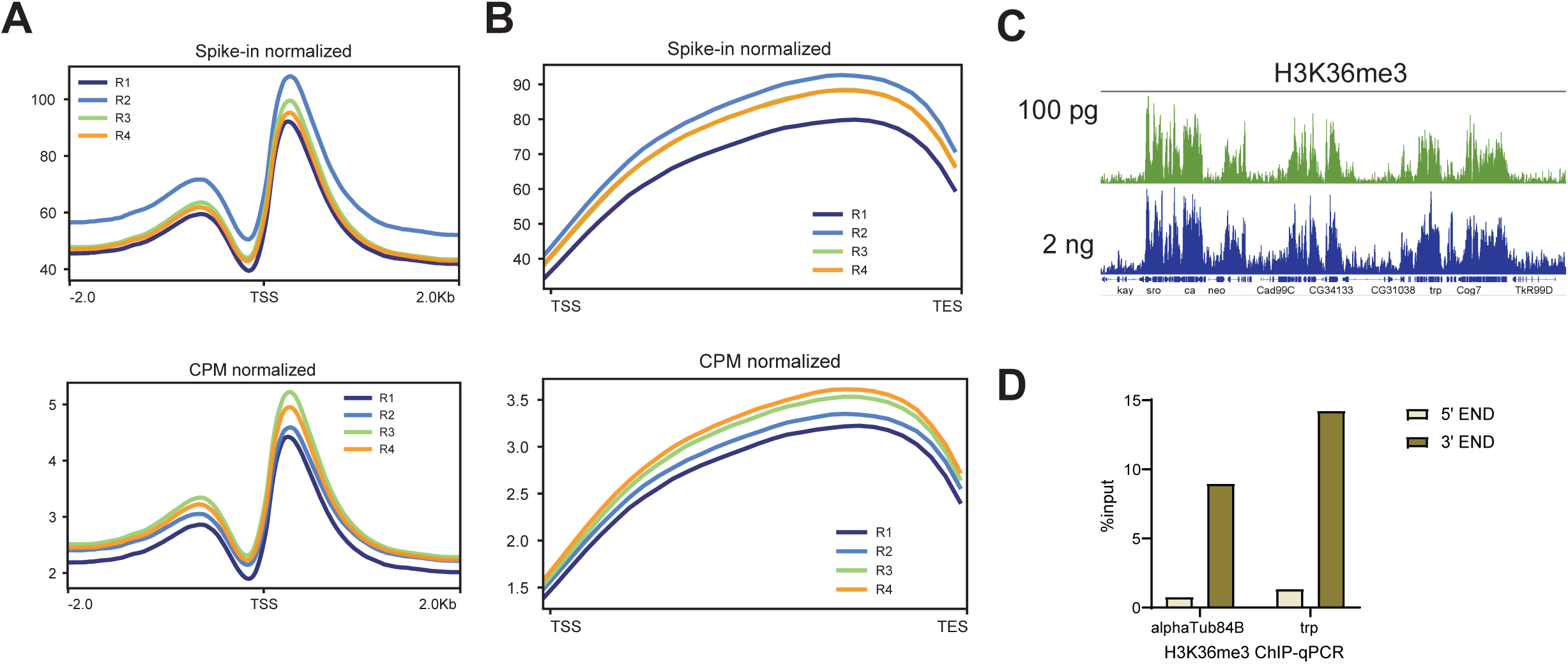
A. H3K4me3 Metaplots of Spike-in (top) and CPM normalized (bottom) data. B. H3K36me3 Metaplots of Spike-in (top) and CPM normalized (bottom) data. C. Genome browser inspection (IGV) of CPM-normalized H3K36me3 signal comparing libraries made with 100 pg or 2 ng of DNA as starting material. D. Bar plot showing H3K36me3 ChIP-qPCR enrichment as percentage chromatin input at the 5’ and 3’ ends of the housekeeping gene *alphaTub84B* and the photoreceptor-specific gene *trp* (n=1).

## References

1. Agrawal, P., Chung, P., Heberlein, U., & Kent, C. (2019). Enabling cell-type-specific behavioral epigenetics in Drosophila: A modified high-yield INTACT method reveals the impact of social environment on the epigenetic landscape in dopaminergic neurons. BMC Biology, 17(1), 30. https://doi.org/10.1186/s12915-019-0646-4

2. Allshire, R. C., & Madhani, H. D. (2018). Ten principles of heterochromatin formation and function. Nature Reviews. Molecular Cell Biology, 19(4), 229–244. https://doi.org/10.1038/nrm.2017.119

3. Ambati, S., Yu, P., McKinney, E. C., Kandasamy, M. K., Hartzell, D., Baile, C. A., & Meagher, R. B. (2016). Adipocyte nuclei captured from VAT and SAT. BMC Obesity, 3. https://doi.org/10.1186/s40608-016-0112-6

4. Amin, N. M., Greco, T. M., Kuchenbrod, L. M., Rigney, M. M., Chung, M.-I., Wallingford, J. B., Cristea, I. M., & Conlon, F. L. (2014). Proteomic profiling of cardiac tissue by isolation of nuclei tagged in specific cell types (INTACT). *Development (Cambridge*, England), 141(4), 962–973. https://doi.org/10.1242/dev.098327

5. Bai, L., & Morozov, A. V. (2010). Gene regulation by nucleosome positioning. Trends in Genetics: TIG, 26(11), 476–483. https://doi.org/10.1016/j.tig.2010.08.003

6. Bailey, M. H., Tokheim, C., Porta-Pardo, E., Sengupta, S., Bertrand, D., Weerasinghe, A., Colaprico, A., Wendl, M. C., Kim, J., Reardon, B., Ng, P. K.-S., Jeong, K. J., Cao, S., Wang, Z., Gao, J., Gao, Q., Wang, F., Liu, E. M., Mularoni, L., … Ding, L. (2018). Comprehensive Characterization of Cancer Driver Genes and Mutations. Cell, 173(2), 371–385.e18. https://doi.org/10.1016/j.cell.2018.02.060

7. Bolger, A. M., Lohse, M., & Usadel, B. (2014). Trimmomatic: A flexible trimmer for Illumina sequence data. Bioinformatics, 30(15), 2114–2120. https://doi.org/10.1093/bioinformatics/btu170

8. Bolus, H., Crocker, K., Boekhoff-Falk, G., & Chtarbanova, S. (2020). Modeling Neurodegenerative Disorders in Drosophila melanogaster. International Journal of Molecular Sciences, 21(9). https://doi.org/10.3390/ijms21093055

9. Bozek, M., Cortini, R., Storti, A. E., Unnerstall, U., Gaul, U., & Gompel, N. (2019). ATAC-seq reveals regional differences in enhancer accessibility during the establishment of spatial coordinates in the Drosophila blastoderm. Genome Research. https://doi.org/10.1101/gr.242362.118

10. Brahma, S., & Henikoff, S. (2020). Epigenome Regulation by Dynamic Nucleosome Unwrapping. Trends in Biochemical Sciences, 45(1), 13–26. https://doi.org/10.1016/j.tibs.2019.09.003

11. Brand, A. H., & Perrimon, N. (1993). Targeted gene expression as a means of altering cell fates and generating dominant phenotypes. Development, 118(2), 401–415.

12. Buenrostro, J. D., Giresi, P. G., Zaba, L. C., Chang, H. Y., & Greenleaf, W. J. (2013). Transposition of native chromatin for fast and sensitive epigenomic profiling of open chromatin, DNA-binding proteins and nucleosome position. Nature Methods, 10(12), 1213–1218. https://doi.org/10.1038/nmeth.2688

13. Chen, K., Hu, Z., Xia, Z., Zhao, D., Li, W., & Tyler, J. K. (2016). The Overlooked Fact: Fundamental Need for Spike-In Control for Virtually All Genome-Wide Analyses. Molecular and Cellular Biology, 36(5), 662–667. https://doi.org/10.1128/MCB.00970-14

14. Chitikova, Z., & Steiner, F. A. (2016). Cell type-specific epigenome profiling using affinity-purified nuclei. Genesis, 54(4), 160–169. https://doi.org/10.1002/dvg.22919

15. Corces, M. R., Trevino, A. E., Hamilton, E. G., Greenside, P. G., Sinnott-Armstrong, N. A., Vesuna, S., Satpathy, A. T., Rubin, A. J., Montine, K. S., Wu, B., Kathiria, A., Cho, S. W., Mumbach, M. R., Carter, A. C., Kasowski, M., Orloff, L. A., Risca, V. I., Kundaje, A., Khavari, P. A., … Chang, H. Y. (2017). An improved ATAC-seq protocol reduces background and enables interrogation of frozen tissues. Nature Methods, 14(10), 959–962. https://doi.org/10.1038/nmeth.4396

16. Davie, K., Jacobs, J., Atkins, M., Potier, D., Christiaens, V., Halder, G., & Aerts, S. (2015). Discovery of Transcription Factors and Regulatory Regions Driving In Vivo Tumor Development by ATAC-seq and FAIRE-seq Open Chromatin Profiling. PLoS Genetics, 11(2). https://doi.org/10.1371/journal.pgen.1004994

17. Deal, R. B., & Henikoff, S. (2010). A simple method for gene expression and chromatin profiling of individual cell types within a tissue. Developmental Cell, 18(6), 1030–1040. https://doi.org/10.1016/j.devcel.2010.05.013

18. Dobin, A., Davis, C. A., Schlesinger, F., Drenkow, J., Zaleski, C., Jha, S., Batut, P., Chaisson, M., & Gingeras, T. R. (2013). STAR: Ultrafast universal RNA-seq aligner. Bioinformatics, 29(1), 15–21. https://doi.org/10.1093/bioinformatics/bts635

19. Edmunds, J. W., Mahadevan, L. C., & Clayton, A. L. (2008). Dynamic histone H3 methylation during gene induction: HYPB/Setd2 mediates all H3K36 trimethylation. The EMBO Journal, 27(2), 406–420. https://doi.org/10.1038/sj.emboj.7601967

20. Göpfert, M. C., & Robert, D. (2001). Turning the key on Drosophila audition. Nature, 411(6840), 908– 908. https://doi.org/10.1038/35082144

21. Graham, P., & Pick, L. (2017). Drosophila as a Model for Diabetes and Diseases of Insulin Resistance. Current Topics in Developmental Biology, 121, 397–419. https://doi.org/10.1016/bs.ctdb.2016.07.011

22. Hales, K. G., Korey, C. A., Larracuente, A. M., & Roberts, D. M. (2015). Genetics on the Fly: A Primer on the Drosophila Model System. Genetics, 201(3), 815–842. https://doi.org/10.1534/genetics.115.183392

23. Hall, H., Ma, J., Shekhar, S., Leon-Salas, W. D., & Weake, V. M. (2018). Blue light induces a neuroprotective gene expression program in Drosophila photoreceptors. BMC Neuroscience, 19(1), 43. https://doi.org/10.1186/s12868-018-0443-y

24. Hall, H., Medina, P., Cooper, D. A., Escobedo, S. E., Rounds, J., Brennan, K. J., Vincent, C., Miura, P., Doerge, R., & Weake, V. M. (2017). Transcriptome profiling of aging Drosophila photoreceptors reveals gene expression trends that correlate with visual senescence. BMC Genomics, 18(1), 894. https://doi.org/10.1186/s12864-017-4304-3

25. Henry, G. L., Davis, F. P., Picard, S., & Eddy, S. R. (2012). Cell type–specific genomics of Drosophila neurons. Nucleic Acids Research, 40(19), 9691–9704. https://doi.org/10.1093/nar/gks671

26. Jones, S. G., Nixon, K. C. J., Chubak, M. C., & Kramer, J. M. (2018). Mushroom Body Specific Transcriptome Analysis Reveals Dynamic Regulation of Learning and Memory Genes After Acquisition of Long-Term Courtship Memory in Drosophila. G3: Genes|Genomes|Genetics, 8(11), 3433–3446. https://doi.org/10.1534/g3.118.200560

27. Kaya-Okur, H. S., Wu, S. J., Codomo, C. A., Pledger, E. S., Bryson, T. D., Henikoff, J. G., Ahmad, K., & Henikoff, S. (2019). CUT&Tag for efficient epigenomic profiling of small samples and single cells. Nature Communications, 10. https://doi.org/10.1038/s41467-019-09982-5

28. Klemm, S. L., Shipony, Z., & Greenleaf, W. J. (2019). Chromatin accessibility and the regulatory epigenome. Nature Reviews Genetics, 20(4), 207–220. https://doi.org/10.1038/s41576-018-0089-8

29. Landt, S. G., Marinov, G. K., Kundaje, A., Kheradpour, P., Pauli, F., Batzoglou, S., Bernstein, B. E., Bickel, P., Brown, J. B., Cayting, P., Chen, Y., DeSalvo, G., Epstein, C., Fisher-Aylor, K. I., Euskirchen, G., Gerstein, M., Gertz, J., Hartemink, A. J., Hoffman, M. M., … Snyder, M. (2012). ChIP-seq guidelines and practices of the ENCODE and modENCODE consortia. Genome Research, 22(9), 1813–1831. https://doi.org/10.1101/gr.136184.111

30. Langmead, B., & Salzberg, S. L. (2012). Fast gapped-read alignment with Bowtie 2. Nature Methods, 9(4), 357–359. https://doi.org/10.1038/nmeth.1923

31. Lardenoije, R., Iatrou, A., Kenis, G., Kompotis, K., Steinbusch, H. W. M., Mastroeni, D., Coleman, P., Lemere, C. A., Hof, P. R., van den Hove, D. L. A., & Rutten, B. P. F. (2015). The epigenetics of aging and neurodegeneration. Progress in Neurobiology, 131, 21–64. https://doi.org/10.1016/j.pneurobio.2015.05.002

32. Li, H., Handsaker, B., Wysoker, A., Fennell, T., Ruan, J., Homer, N., Marth, G., Abecasis, G., & Durbin, R. (2009). The Sequence Alignment/Map format and SAMtools. Bioinformatics, 25(16), 2078–2079. https://doi.org/10.1093/bioinformatics/btp352

33. Liao, Y., Smyth, G. K., & Shi, W. (2013). The Subread aligner: Fast, accurate and scalable read mapping by seed-and-vote. Nucleic Acids Research, 41(10), e108. https://doi.org/10.1093/nar/gkt214

34. Ma, J., Brennan, K. J., D’Aloia, M. R., Pascuzzi, P. E., & Weake, V. M. (2016). Transcriptome Profiling Identifies Multiplexin as a Target of SAGA Deubiquitinase Activity in Glia Required for Precise Axon Guidance During Drosophila Visual Development. G3: Genes|Genomes|Genetics, 6(·), 2435–2445. https://doi.org/10.1534/g3.116.031310

35. Ma, J., & Weake, V. M. (2014). Affinity-based Isolation of Tagged Nuclei from Drosophila Tissues for Gene Expression Analysis. JoVE (Journal of Visualized Experiments), 85, e51418. https://doi.org/10.3791/51418

36. Maher, K. A., Bajic, M., Kajala, K., Reynoso, M., Pauluzzi, G., West, D. A., Zumstein, K., Woodhouse, M., Bubb, K., Dorrity, M. W., Queitsch, C., Bailey-Serres, J., Sinha, N., Brady, S. M., & Deal, R. B. (2018). Profiling of Accessible Chromatin Regions across Multiple Plant Species and Cell Types Reveals Common Gene Regulatory Principles and New Control Modules. The Plant Cell, 30(1), 15–36. https://doi.org/10.1105/tpc.17.00581

37. Mollereau, B., Wernet, M. F., Beaufils, P., Killian, D., Pichaud, F., Kühnlein, R., & Desplan, C. (2000). A green fluorescent protein enhancer trap screen in Drosophila photoreceptor cells. Mechanisms of Development, 93(1), 151–160. https://doi.org/10.1016/S0925-4773(00)00287-2

38. Orlando, D. A., Chen, M. W., Brown, V. E., Solanki, S., Choi, Y. J., Olson, E. R., Fritz, C. C., Bradner, J. E., & Guenther, M. G. (2014). Quantitative ChIP-Seq Normalization Reveals Global Modulation of the Epigenome. Cell Reports, 9(3), 1163–1170. https://doi.org/10.1016/j.celrep.2014.10.018

39. Piper, M. D. W., & Partridge, L. (2018). Drosophila as a model for ageing. Biochimica et Biophysica Acta (BBA) - Molecular Basis of Disease, 1864(9, Part A), 2707–2717. https://doi.org/10.1016/j.bbadis.2017.09.016

40. Potter, C. J., Tasic, B., Russler, E. V., Liang, L., & Luo, L. (2010). The Q system: A repressible binary system for transgene expression, lineage tracing, and mosaic analysis. Cell, 141(3), 536–548. https://doi.org/10.1016/j.cell.2010.02.025

41. Ramírez, F., Dündar, F., Diehl, S., Grüning, B. A., & Manke, T. (2014). deepTools: A flexible platform for exploring deep-sequencing data. Nucleic Acids Research, 42(Web Server issue), W187–W191. https://doi.org/10.1093/nar/gku365

42. Sijacic, P., Bajic, M., McKinney, E. C., Meagher, R. B., & Deal, R. B. (2018). Changes in chromatin accessibility between Arabidopsis stem cells and mesophyll cells illuminate cell type-specific transcription factor networks. The Plant Journal: For Cell and Molecular Biology, 94(2), 215–231. https://doi.org/10.1111/tpj.13882

43. Skene, P. J., & Henikoff, S. (2017). An efficient targeted nuclease strategy for high-resolution mapping of DNA binding sites. ELife, 6, e21856. https://doi.org/10.7554/eLife.21856

44. Slankster, E., Kollala, S., Baria, D., Dailey-Krempel, B., Jain, R., Odell, S. R., & Mathew, D. (2020). Mechanism underlying starvation-dependent modulation of olfactory behavior in Drosophila larva. Scientific Reports, 10(1), 3119. https://doi.org/10.1038/s41598-020-60098-z

45. Stadler, J., & Richly, H. (2017). Regulation of DNA Repair Mechanisms: How the Chromatin Environment Regulates the DNA Damage Response. International Journal of Molecular Sciences, 18(8). https://doi.org/10.3390/ijms18081715

46. Stark, W. S., & Thomas, C. F. (2004). Microscopy of multiple visual receptor types in Drosophila. Molecular Vision, 10, 943–955.

47. Ugur, B., Chen, K., & Bellen, H. J. (2016). Drosophila tools and assays for the study of human diseases. Disease Models & Mechanisms, 9(3), 235–244. https://doi.org/10.1242/dmm.023762

48. Wingett, S. W., & Andrews, S. (2018). FastQ Screen: A tool for multi-genome mapping and quality control. F1000Research, 7, 1338. https://doi.org/10.12688/f1000research.15931.2

49. Yu, G., Wang, L.-G., Han, Y., & He, Q.-Y. (2012). clusterProfiler: An R Package for Comparing Biological Themes Among Gene Clusters. OMICS : A Journal of Integrative Biology, 16(5), 284–287. https://doi.org/10.1089/omi.2011.0118

50. Yu, G., Wang, L.-G., & He, Q.-Y. (2015). ChIPseeker: An R/Bioconductor package for ChIP peak annotation, comparison and visualization. Bioinformatics, 31(14), 2382–2383. https://doi.org/10.1093/bioinformatics/btv145

51. Zhang, Y., Liu, T., Meyer, C. A., Eeckhoute, J., Johnson, D. S., Bernstein, B. E., Nusbaum, C., Myers, R. M., Brown, M., Li, W., & Liu, X. S. (2008). Model-based analysis of ChIP-Seq (MACS). Genome Biology, 9(9), R137. https://doi.org/10.1186/gb-2008-9-9-r137

